# Simultaneous, cortex-wide and cellular-resolution neuronal population dynamics reveal an unbounded scaling of dimensionality with neuron number

**DOI:** 10.1101/2024.01.15.575721

**Authors:** Jason Manley, Jeffrey Demas, Hyewon Kim, Francisca Martínez Traub, Alipasha Vaziri

## Abstract

The brain’s remarkable properties arise from collective activity of millions of neurons. Widespread application of dimensionality reduction to multi-neuron recordings implies that neural dynamics can be approximated by low-dimensional “latent” signals reflecting neural computations. However, what would be the biological utility of such a redundant and metabolically costly encoding scheme and what is the appropriate resolution and scale of neural recording to understand brain function? Imaging the activity of one million neurons at cellular resolution and near-simultaneously across mouse cortex, we demonstrate an unbounded scaling of dimensionality with neuron number. While half of the neural variance lies within sixteen behavior-related dimensions, we find this unbounded scaling of dimensionality to correspond to an ever-increasing number of internal variables without immediate behavioral correlates. The activity patterns underlying these higher dimensions are fine-grained and cortex-wide, highlighting that large-scale recording is required to uncover the full neural substrates of internal and potentially cognitive processes.

## INTRODUCTION

Individual neurons in the brain are not independent processors, but densely interconnected and interdependent units that collectively enable computations underlying the execution of adaptive and goal-directed behavior. As such, it has long been argued that unraveling the dynamics of neural computation will require the ability to monitor the firing patterns of large neuronal populations distributed throughout the brain (Abbott and Dayan, 1999; Perkel et al., 1967).

However, until the last decade, simultaneous recordings were limited to only a few cells at a time (Nicolelis, 2007; Stevenson and Kording, 2011), even in the simplest model systems (Zheng et al., 2012). As a result, some of the key paradigms and conceptual frameworks in neuroscience have been based on measurable properties of single neurons, and corresponding theoretical frameworks for neural computation have aimed to describe the responses of individual cells and infer how they could arise from the large-scale but often unobservable neural circuitry within which they are embedded (Barlow, 1961; Fairhall et al., 2001; Parker and Newsome, 1998; Shadlen and Newsome, 1998; Vreeswijk and Sompolinsky, 1996).

With the advent of large-scale neuronal recording, it is now possible to monitor the *in vivo* activity of increasingly larger ensembles of neurons simultaneously (Grewe and Helmchen, 2009; Tsai et al., 2015; Cai et al., 2016; Carrillo-Reid et al., 2016; Prevedel et al., 2016; Sofroniew et al., 2016; Zong et al., 2017; Nöbauer et al., 2017; Skocek et al., 2017; Steinmetz et al., 2018; Weisenburger and Vaziri, 2018; Weisenburger et al., 2019; Rumyantsev et al., 2020; Demas et al., 2021; Yu et al., 2021; Kim and Schnitzer, 2022; Urai et al., 2022; Zong et al., 2022) while animals are engaged in behavior, in some cases up to the level of whole-brains (Schrödel et al., 2013; Ahrens et al., 2013; Prevedel et al., 2014; Nguyen et al., 2016). Given the ever-growing number of recorded units, dimensionality reduction has emerged as a powerful tool for visualization and interpretation of neural population dynamics, as well as for quantitative analysis of the features they encode (Cunningham and Yu, 2014; Pang et al., 2016). The basic underlying assumption in dimensionality reduction is that the *N* measured variables covary according to a smaller set of variables, often called “latent” variables, as they are not measured directly but inferred from the structure of the observed variables. In geometric terms, the full *N*-dimensional state space is thought to be constrained to a lower-dimensional manifold, and dimensionality reduction methods aim to infer this manifold structure and extract the encoded latent variables.

In practice, this view has been supported by empirical evidence showing that under a number of experimental conditions the estimated dimensionality of neural dynamics is consistently much lower than the full state space (Gao and Ganguli, 2015) and that relatively few latent signals often capture a large proportion of the relevant behavior or stimulus-related activity (Gallego et al., 2017; MacDowell and Buschman, 2020; Seung and Lee, 2000). This distributed and redundant coding scheme across many neurons has been proposed to provide robustness and resiliency to noise (Montijn et al., 2016; Shadlen and Newsome, 1998; Zylberberg et al., 2017, 2016), yet it can significantly reduce the information encoding capacity of the neural population (Kafashan et al., 2021; Moreno-Bote et al., 2014; Rumyantsev et al., 2020). At the same time, the ubiquity of these low-dimensional, redundant neural codes suggests that it may be possible to recover the relevant latent dynamics by measuring just a subset of the full neural population.

However, recent studies combining large-scale recording and novel dimensionality reduction techniques suggest that the neural dimensionality of a population, defined as the minimum number of continuous variables needed to parameterize the neural dynamics (Jazayeri and Ostojic, 2021), is higher than previously appreciated. For example, it has been shown that signals in a region of visual cortex can contain over 100 significant latent variables, many of which encode for ongoing animal behavior as opposed to solely visual stimuli (Stringer et al., 2019b).

Other studies have suggested that the observed neural dimensionality is inherited from the task or the stimulus structure. In fact, it was shown that the geometry within visual cortex is related to the input stimulus dimensionality, such that the neural dimensionality is maximized while maintaining smoothness of the neural code (Stringer et al., 2019a). Coding smoothness enforces that similar stimuli evoke similar population responses, preventing a fractal and undifferentiable geometry where small changes in stimulus evoke dramatic changes in the neural population activity, allowing for flexible coding that maintains robustness and generalization. High-dimensional representations have also been observed in more cognitive tasks (Bernardi et al., 2020; Parthasarathy et al., 2017; Rigotti et al., 2013), where it has been argued that mixed selectivity in neurons increases the neuronal dimensionality and is critical for cognition (Fusi et al., 2016). Together these results suggest that neural circuits encode as many relevant features as possible – perhaps including unexpected features, such as the case of behavior encoding across sensory cortex – while preserving a correlation structure that balances robustness, efficiency, and flexibility.

Despite the increasing evidence for high-dimensional neural geometry, an open fundamental question is how neuronal dimensionality would scale with neuron number, which has only been studied within a limited neuronal population size (Williamson et al., 2016). If the neural population dynamics associated with a given task or behavior can be truly represented by a low-dimensional system and a sufficiently large number of neurons are recorded, then the measured dimensionality would be expected to exhibit a bounded scaling, i.e. the measured dimensionality would plateau as more neurons are recorded (Figure 1A, orange line). This view, also referred to as the “strong principle” of dimensionality reduction (Humphries, 2020), would suggest that neural circuits are tuned to encode information redundantly in their lower-dimensional activity patterns and that the utilized recording technology has sampled the population sufficiently to reveal its underlying manifold structure. On the other hand, the observation of unbounded scaling during any type of behavior or task, such that dimensionality is ever-increasing with neuron number up to the level of the entire recorded neuronal population (Figure 1A, blue line), would suggest that while dimensionality reduction can still serve as a useful interpretive technique in certain paradigms, the utilized recording technology still lies in a regime of undersampling. In that case, full observation of brain state dynamics would require a larger sampling of the neuronal population, potentially even whole-brain recording. Importantly, these two scalings underly highly divergent neural coding schemes, which imply fundamentally different structural and functional properties of the underlying neural circuitry and would require distinct experimental and analytical methods to infer the full set of latent encoded variables.

**Figure 1.**
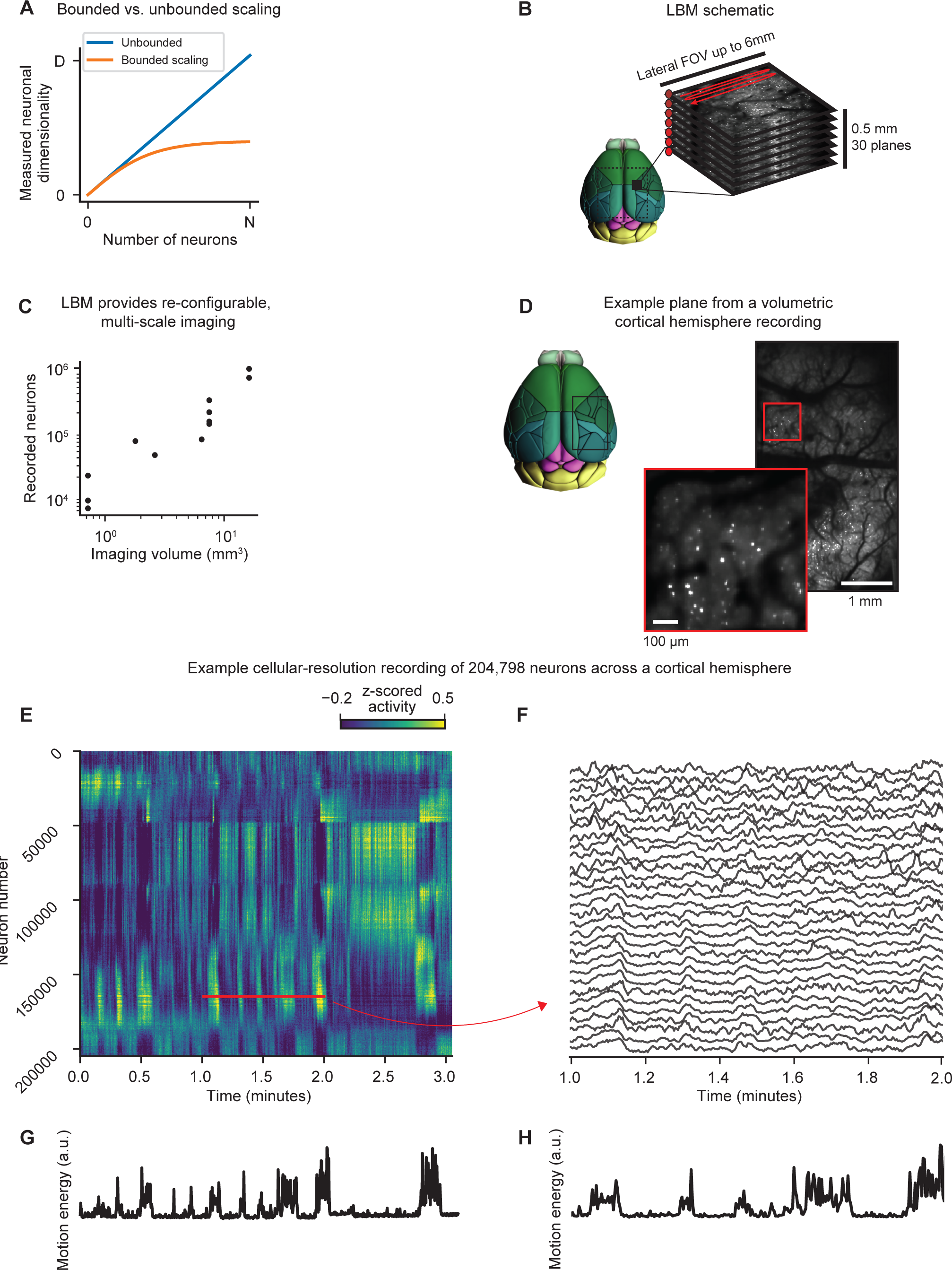
Light Beads Microscopy (LBM) enables large-scale, volumetric recording of neuronal activity across cortex at cellular resolution. **A.** A schematic representation of two possible scenarios: bounded versus unbounded scaling of the measured neuronal dimensionality as a function of number of recorded neurons. **B.** A schematic of the LBM imaging setup. A column of 30 successive axially-separated and temporally-delayed foci or “light beads” excite individual planes separated by ∼16 µm over a total axial range of 0.5 mm within the mouse cortex. Fluorescence from a single column is captured within 200 ns. The column of light beads is then scanned across a lateral field of view (FOV) with a maximum size of 6 mm (represented by the dashed area on the mouse brain rendering reproduced by Brain Explorer 2 (Lau et al., 2008)), thereby enabling cellular resolution, volumetric recording of neuronal activity at multi-Hertz rate. **C.** The number of recorded neurons is proportional to the size of the imaged volume, ranging from 6,519 to 970,546 neurons in n=12 recordings. **D.** An example single hemisphere recoding. Imaging was done within a volume of ∼3.0 x 5.0 x 0.5 mm imaged at ∼5 µm lateral sampling for 1 hour at 4.7 Hz, resulting in 204,798 identified neurons. Top left: the approximate imaging location denoted by a black box. The recording volume covers portions of the primary motor, primary somatosensory, posterior parietal, retrosplenial, primary visual, anteromedial visual, and posteromedial visual cortices. Right: The standard deviation projection of a plane 183 µm below the cortical surface and corresponding zoom in. Scale bar: 1 mm. Inset scale bar: 100 µm. **E.** Z-scored heatmap of neural activity from a 3-minute portion of the single hemisphere recording in panel D. The neurons are sorted utilizing the rastermap algorithm (Stringer et al. 2019b), which sorts neurons based on similarity of their temporal activity patterns. **F.** 30 representative examples of individual neuronal traces from the red highlighted region in E. **G** and **H.** Simultaneously recorded behavior activity for the neural dynamics shown in panels E and F, respectively. The behavior is coarsely quantified here as the total motion energy, defined as the sum of the absolute difference in pixel values between consecutive frames.

Here, we hypothesized that previous observations of low-dimensional neuronal activity were the result of limitations in either the recorded population size or in accurately identifying all signal-carrying dimensions. To investigate this, we systematically investigated how the measured neural dimensionality scales with population size by recording the dynamics of up to one million neurons distributed across different depths and regions of mouse dorsal cortex using light beads microscopy (LBM) (Demas et al., 2021) during spontaneous and uninstructed behavior in head-fixed mice. Utilizing a recent approach to identify dimensions of reliable neural variance, we demonstrated that the measured dimensionality scales according to an unbounded power law with neuron number, suggesting that even at this scale of recording our technologies are still undersampling the full state space of neural dynamics for spontaneous cortical activity.

Consistent with previous reports (Kauvar et al., 2020; Musall et al., 2019; Salkoff et al., 2020; Stringer et al., 2019b), we identified a low-dimensional encoding of the animal’s behavior at each timepoint by approximately sixteen of the largest dimensions, which collectively accounted for roughly half of the observed neural variance. Our large-scale, cellular resolution, and volumetric recordings revealed that each of these behavior-related signals was encoded by multiple spatially localized clusters of covarying neurons distributed across both cortical hemispheres. Taken together, our results suggest that these initial largest dimensions form a cortex-wide (and potentially brain-wide) network in which behavioral information is broadcast in a global manner by spatially clustered ensembles of functionally interconnected neurons.

On the other hand, given that so few neural dimensions were related to the animal’s motor activity, the observed increase in reliable dimensionality as a function of neuron number implied that the higher reliable dimensions represented neural modes without any immediate relationship to behavior. Nevertheless, we found that these higher dimensional components indeed exhibited a temporal structure that is distinct from noise and spanned a continuum of timescales from seconds to the limit of the temporal resolution in our recordings. Also, given the lack of any sensory inputs to the animals, we reason that these dimensions form a high-dimensional geometry representing dynamics underlying purely internal neural computations that may enable adaptive behavior across a wide range of timescales, however the specific nature and function of the information these signals reliably encode remains to be identified. Enabled by the large-scale neuronal recording capability of LBM, our results have revealed the high-dimensional geometry of spontaneous cortical dynamics and demonstrate that application of low-rank dimensionality reduction to these data would lead to loss of information, as well as systematic biases in the observed spatial profiles, timescales, and encoded features.

## RESULTS

To investigate the geometry and dimensionality of spontaneous neuronal population dynamics across different depths and regions of the dorsal cortex of the mouse, we used LBM which allowed for volumetric, single cell resolution recording of calcium dynamics from populations of up to one million neurons (Figure 1B). In LBM, a recent volumetric calcium imaging technique, the fluorescent activity of neurons within a 3D imaging volume is captured using a column of 30 axially separated and temporally distinct two-photon excitation spots or “light beads”. These light beads capture neural activity densely along the axial range in rapid succession, providing a spatiotemporally optimal acquisition that is limited only by the fluorescence lifetime of the genetically encodable calcium indicator. The column of light beads is subsequently scanned across the lateral field of view (FOV) to sample a 3D mesoscale imaging volume at cellular resolution (Video 1). LBM’s versatility allows for recordings ranging from 1.2 x 1.2 x 0.5 mm at 10 Hz and 2 µm lateral voxel spacing to 5.4 x 6.0 x 0.5 mm at 2.2 Hz and 5 µm spacing (see examples in Figure S1), yielding populations of 6,519 to 970,546 near-simultaneously recorded neurons, respectively (Figure 1C). We imaged transgenic mice expressing the genetically encoded calcium indicator GCaMP6s or GCaMP6f (Daigle et al., 2018) in glutamatergic neurons. The use of an 8 mm diameter chronic cranial window implantations (see STAR Methods for details) allowed for optical access to the majority of the dorsal cortical volume, including the visual, somatosensory, posterior parietal, retrosplenial, and motor cortices across both cortical hemispheres (Figure 1D). Neural activity across these cortical regions was imaged for one hour while animals were free to move on a linear treadmill and exhibited epochs of various spontaneous and uninstructed behaviors such as whisking, grooming, and locomotion (Video 2).

It has been shown that even in expertly-trained animals performing decision making tasks, mesoscale cortical activity appears dominated by task-irrelevant, uninstructed behavioral signals (Kauvar et al., 2020; Musall et al., 2019; Salkoff et al., 2020), however the exact function and dimensionality of this widespread and redundant behavioral encoding is unclear as it has not been studied at cellular resolution across such large FOVs simultaneously. Additionally, we chose this spontaneous arrangement given that it provides minimal external shaping of the neural dimensionality without a task or stimulus structure. Generally, the obtained neural recordings exhibited different spatiotemporal activity patterns at multiple scales (Figures 1E-F). Many of these activity patterns were correlated with the animal’s simultaneous epochs of motor behavior, as quantified using motion energy, which is defined as the total absolute pixel-wise difference between subsequent behavioral video frames (Figures 1G-H). Together, these neurobehavioral data represent the largest available cellular-resolution and near-simultaneous recordings of neuronal population dynamics during behavior, providing a unique opportunity to assess the geometry and spatiotemporal structure of cortex-wide neuronal dynamics.

### Reliable dimensionality of cortex-wide neuronal dynamics during spontaneous behavior exhibits unbounded power law scaling with neuron number

To date, technical limitations on the ability to record large-scale or brain-wide activity of neuronal populations at cellular resolution and to accurately estimate their underlying dimensionality have been the prohibitive factor to determine whether their dynamics are confined to a low-dimensional manifold. Given recent observations of high-dimensionality neural geometry using large-scale imaging techniques, we hypothesized that the observed neuronal dimensionality would only continue to grow as neuron number surpassed previous recordings on the order of 10,000 neurons. To evaluate this conjecture, we utilized the large-scale recording capabilities of LBM to characterize the structure and dimensionality of cortex-wide neural activity by identifying dimensions of neural covariance that encode underlying latent signals shared across neurons. We used shared variance component analysis (SVCA) (Stringer et al., 2019b) as an alternative to traditional dimensionality reduction techniques to identify globally shared components of variation in neuronal population dynamics. A key feature of SVCA is that it allows distinguishing between dimensions that truly contain reliable signal versus those that represent noise and overfitting to the training set due to finite sampling.

SVCA first divides the neurons into two subsets, which are chosen by segmenting the lateral FOV into 250 µm squares and collating every other square in a checkerboard pattern, thus preventing any axial crosstalk between the two sets (Figure 2A(i)). Similarly, their timepoints are divided into equal training and testing sets for cross-validation by chunking the full recording into 72 second intervals and randomly assigning each chunk to either the training or testing timepoints. Utilizing the training timepoints, SVCA identifies the planar dimensions of each neural set’s activity that maximally covary, called the shared variance components (SVCs, see Figure 2A(ii) and STAR Methods for details). In order to determine which of these neural SVCs contain meaningful signal, the reliability of each SVC is measured by its covariance between the two neural sets on the held-out testing timepoints (Figure 2A(iii)). This reliability measure allows for identification of all dimensions containing robust dynamics, as opposed to those representing noise or otherwise unreliable features due to overfitting to the finite number of training timepoints. This approach has at least two advantages compared to traditional dimensionality reduction techniques, such as PCA that lack cross-validation required to quantify the latent signals’ reliability. First, it allows for a separation between the quality of a latent signal – which is related to its potential biological relevance – and the amount of variance it explains.

**Figure 2.**
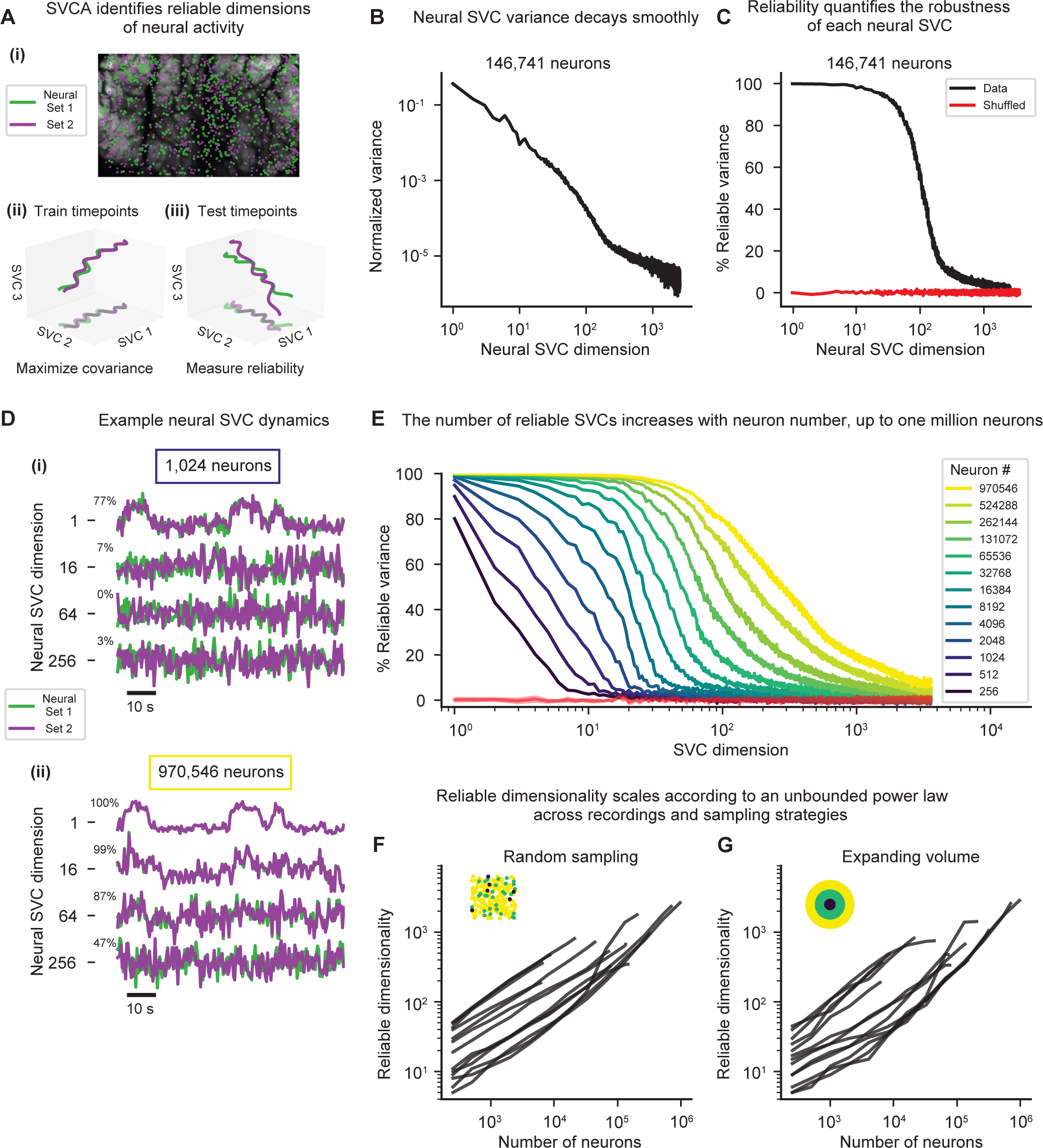
Shared variance component analysis (SVCA) reveals unbounded scaling of reliable neural dimensionality. **A.** Schematics of SVCA: (i): SVCA splits the neurons into two sets (green and purple), which are shown by their lateral location over a corresponding cortical hemisphere image. (ii): The training timepoints are used to identify the shared variance components (SVCs), which are the maximally covarying projections of each neural set. (iii): To quantify whether these SVCs contain biological signal versus noise, the reliability of each neural SVC is quantified using the covariance of the two sets’ projections on held-out testing timepoints. **B.** The normalized variance spectra across neural SVC dimensions decays smoothly. Data from an example hemisphere recording containing 146,741 neurons. **C.** The percentage of reliable variance quantifies the robustness of each SVC, which eventually decreases at higher SVCs. The data for the same population as in panel B is shown in black, whereas temporally shuffled data is shown in red, where each neuron’s timepoints are split into portions of two seconds in duration and then randomly permuted. **D.** Visualization of the reliability of individual SVCs as a function of neuronal population size. The reliability of each SVC can be visualized by comparing the two neural sets’ SVC projections. SVC signals from a random sampling of (i): 1,024 neurons and (ii): 970,546 neurons are shown for the bi-hemisphere recording analyzed in panel E. The percentage of reliable variance in each SVC is shown on the left of the corresponding traces. **E.** The dimensionality of the neuronal population as quantified by reliable SVCs exhibits unbounded scaling as a function of the number of recorded neurons. Colored traces show different size (256 – 970,546) neural populations randomly sampled from a bi-hemispheric recorded volume with 970,546 neurons, with each trace indicating the mean percentage of reliable variance in each SVC over n=10 iterations of random sampling. The red shaded line indicates the mean ± 95% CI for temporally shuffled data (see STAR Methods for details) from 970,546 neurons. **F** and **G.** The reliable neural dimensionality exhibits unbounded scaling with neuron number across different neuronal sampling strategies. The reliable neural dimensionality is defined as the number of neural SVC dimensions with a reliable variance percentage greater than four standard deviations above the mean of corresponding temporally shuffled data. Each line indicates the mean over n=10 iterations of random sampling in a single mouse recording, ranging from 1.2 x 1.2 x 0.5 mm to 5.4 x 6 x 0.5 mm FOVs. The reliable dimensionality exhibits unbounded power law scaling as a function of the number of sampled neurons. In F, neurons are sampled randomly from the entire volume (depicted in the pictogram on the top left, the colors which correspond to the legend in panel E), while in G, neurons are sampled in order of their distance from the center of the volume.

This is unlike PCA in which some ad hoc chosen amount of cumulative variance is generally used to justify the dimensionality of the system as the number of components representing that variance. Second, given that the variance within each component is generally smoothly decaying with component number (Figure 2B) and it is difficult to estimate *a priori* what fraction of the data contains neural signal versus noise, the above choice of the variance threshold can be an arbitrary decision. Thus, in contrast to PCA and similar methods, SVCA’s reliability approach provides a data-driven estimate of the latent dimensionality as the number of significantly reliable SVCs compared to the noise floor or a shuffling test.

Applying SVCA to a cortical hemisphere recording from a volume of 3 x 5 x 0.5 mm at 4.7 Hz containing 146,741 total neurons, we found that more than 1,000 dimensions exhibited greater reliability than temporally shuffled data (Figure 2C), in which each neuron’s timeseries was chunked into two second time intervals and then randomly permuted (see STAR Methods for details). Next, to quantify how the number of reliable SVCs behaved as a function of the number of sampled neurons, we performed SVCA on various subsets of neurons sampled randomly from across the imaging volume. While visual inspection of the projections of both neural sets on the held-out testing timepoints indicated that at low neuron numbers (e.g. 1024 neurons) only a few dimensions exhibited reliable covariations, many more components exhibited strong covariation as the number of sampled neurons increased up to our largest recordings of 970,546 neurons (Figure 2D and S2A). This was unlike what was found in data that was temporally shuffled as described previously (Figure S2B). For example, we found that while small populations of only hundreds of neurons contained on the order of ten reliable SVCs, our largest sampling of nearly one million neurons contained thousands of SVCs with greater reliability than in shuffled data (2,664 ± 104 reliable dimensions, mean ± 95% confidence interval across n=10 neuronal samplings, Figure 2E). In this context we defined reliable dimensionality as the number of SVCs in which the percentage of reliable variance was greater than four standard deviations above the mean of a temporally shuffled dataset. This analysis revealed an unbounded power law scaling of reliable dimensionality with the number of sampled neurons (Figure 2F), with a power law exponent of a-0.67 ± 0.04 (mean ± 95% confidence interval across n=12 recordings). This power law exponent indicates the increase in dimensionality is sublinear yet monotonically increasing with neuron number. Moreover, we found this unbounded scaling of reliable dimensionality was consistent across other arbitrary choices for a minimum reliable variance threshold (Figure S2C).

Taken together, our data highlight the high-dimensional geometry of spontaneous cortical dynamics and suggest that the measured dimensionality will only continue to grow with increasing recording capacity. In fact, we believe these estimates likely represent only a lower bound on the true dimensionality of the cortical population activity. First, we are sampling at most an estimated 10% of cortical neurons, and our unbounded scaling law suggests the dimensionality will continue to grow with neuron number. Secondly, our largest recordings have been performed at lower volume rates due to LBM’s inherent tradeoffs between speed, volume size, and sampling density. In this context, post-hoc temporal down-sampling analyses (see STAR Methods for details) on our 10 Hz recordings revealed that the reliable dimensionality drops significantly in the low single Hz volume acquisition rate regime, with about 40% fewer reliable dimensions at 2 Hz volume rate (Figure S2D). Finally, we recorded neural dynamics for an hour, which resulted in many recordings that contained significantly fewer measured timepoints than total number of neurons. While this could in principle limit the number of measurable reliable SVCs due to insufficient sampling of the neural activity patterns, in our particular case we found that the measured reliable dimensionality saturated within this one-hour recording duration for all but our largest bi-hemispheric recordings (Figure S2E), suggesting our recording duration was sufficient to robustly capture the full reliable dimensionality of neural dynamics within nearly all of our experiments. Nevertheless, despite these trade-offs, we ultimately observed the largest reliable dimensionality in our nearly one million neuron bi-hemispheric recordings taken at 2 Hz volume rate (Figure 2E).

Notably, our observed unbounded scaling of the reliable dimensionality of the neural population activity also highlights the limitation of traditional variance-based approaches that do not quantify the reliability of the neural dimensions via cross-validation. We found that the variance spectra as a function of SVC dimension exhibited only minor changes with neuron number (Figure S2F), suggesting that the amount of variance within each SVC does not necessarily correlate with its reliability. Accordingly, we found that the number of principal components (PCs) required to explain some *ad hoc* percentage of the total variance exhibited bounded scaling as a function of neuron number (Figure S2G). This discrepancy is due to the fact that a component may explain only a small fraction of the variance, but it can still be found to contain reliable information as long as the number of sampled neurons is sufficiently large. In other words, the minimum variance threshold required to separate reliable dimensions from noise depends on the number of sampled neurons.

Finally, given that our above observations (Figure 2D-F) were based on random sampling from across the entire imaged volume, which contained multiple cortical regions or hemispheres, we asked whether our observed scaling of reliable dimensionality depended on the spatial sampling pattern of neurons. To address this question, we sampled from an expanding FOV, in which neurons were sampled in order of their distance from the center of the volume. Interestingly, we found our observed scaling law was independent of the sampling strategy; the expanding FOV sampling exhibited a consistent power law exponent of a-0.71 ± 0.06 (Figure 2G), not significantly different from random sampling (p=0.21, two-sided t-test). We further corroborated this observation by confirming that this scaling was also conserved for different neuronal sampling densities by sampling the same neuron number from different volume sizes (Figure S2H).

The above observations demonstrate that the dimensionality of spontaneous cortical dynamics is primarily dependent on the neuron number, as opposed to their spatial distribution across the volume or among cortical regions. This suggests that many of the variables encoded in the latent SVCs may be widely and potentially homogeneously distributed across the cortex, consistent with previous findings based on widefield imaging lacking cellular resolution (Kauvar et al., 2020; Musall et al., 2019; Wekselblatt et al., 2016). Taken together, our results demonstrate that this unbounded power law scaling is consistent across recordings, experimental configurations, and sampling strategies, suggesting that cortical population dynamics indeed encode a very high-dimensional and distributed set of latent signals that cannot be detected without simultaneous observation of the activity of very large numbers of neurons at cellular resolution.

### Low-dimensional encoding of motor behavior implies that the majority of neural SVCs encode purely internal variables

Linking such inferred latent dimensions to sensory inputs, motor actions, and cognitive processes is key to understanding neural circuit computations. Thus, we next sought to identify how behavior-related information is distributed across the neural SVCs. Specifically, we asked whether the observed unbounded scaling of the reliable dimensionality of cortical dynamics represents an increase in the number of encoded behavior-related features or an increase in the number internal features, defined as neural signals that lack a motor or sensory correlate and which could be related to cognitive processes or internal states such as motivational drives.

To differentiate between these two alternatives, we captured the animals’ spontaneous and uninstructed epochs of behavior using a high-speed camera pointed at the animals’ face and body, which allowed for tracking of the majority of the animal postures over time (Video 2). After closer examination and initial analysis of the recordings, we found the animals’ facial movements captured most of the behavioral dynamics that were correlated with the observed neural activity. Thus, we quantified the spontaneous and uninstructed behavior of each mouse by computing the first 500 PCs of facial motion energy between frames (Figure 3A-B, see STAR Methods for details), which we refer to as behavior PCs. The observed behavior appeared multi-dimensional and consistent across mice and imaging sessions (Figure S3A), with 256 PCs explaining 92 ± 3% of the captured motion energy variance. Additionally, the behavior PC activity exhibited various clusters of behavioral motifs (Figure S3B-C), providing a useful moment-to-moment description of spontaneous behavior.

**Figure 3.**
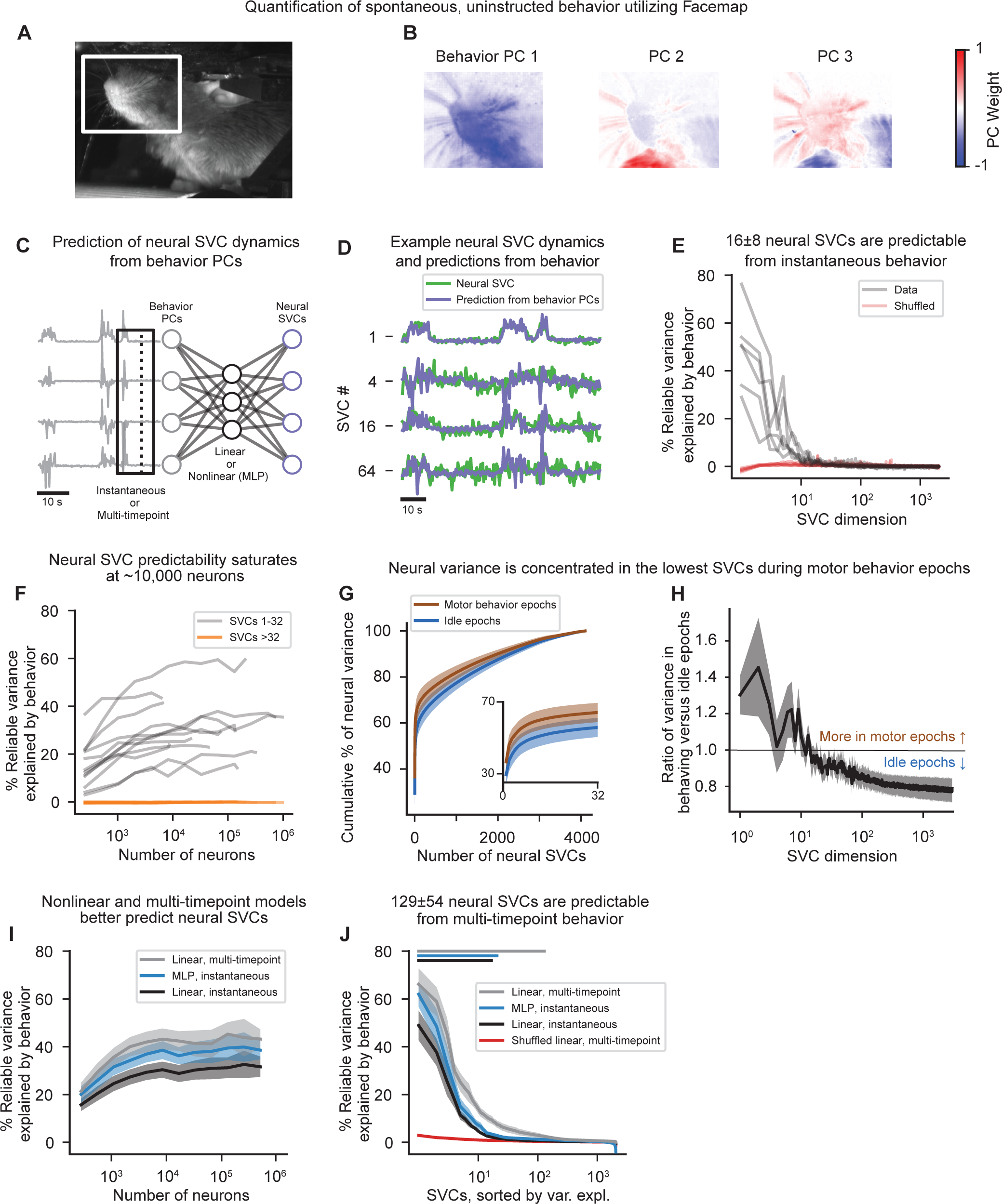
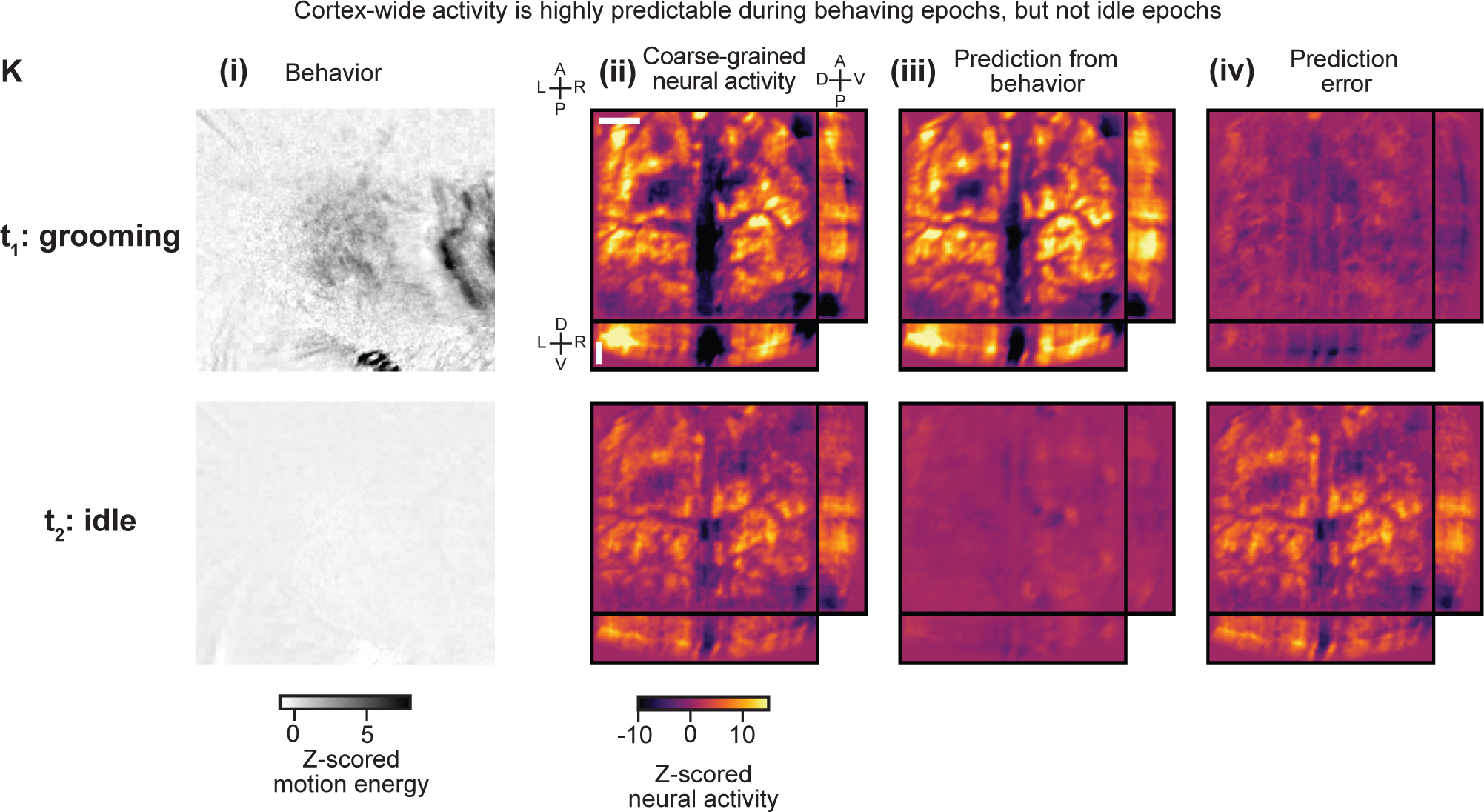
Low-dimensional encoding of behavior implies that the majority of neural SVCs encode purely internal variables. **A.** Example behavior videography image, with a box denoting the area in which facial motion energy is monitored. **B.** The top three behavior principal components (PCs) for an example mouse, identified using the Facemap algorithm. **C.** Schematics of the prediction of neural SVCs from the behavior PCs. Neural SVCs are predicted from the first 500 behavior PCs using either a linear (reduced-rank regression) or nonlinear (multilayer perceptron (MLP), with one hidden layer and ReLU activation) regressor. While all of the models predict a single timepoint of neural SVC activity, the behavior PC inputs are either: instantaneous, such that a single timepoint of behavior PC is provided; or multi-timepoint. In multi-timepoint models, a variable duration behavioral time window around each instantaneous timepoint is utilized, such that time-lagged copies of the behavior PCs are provided as additional inputs to the model to predict a single timepoint of neural SVC activity. **D.** Example z-scored SVC timeseries (green) and their linear, instantaneous predictions from behavior PCs (purple) using held-out testing timepoints. While the initial SVCs appear highly predictable from behavior, the higher order SVCs (e.g., SVC 64) do not. **E.** Only the first 16 ± 8 neural SVCs are predictable from instantaneous behavior. Using the linear, instantaneous reduced-rank regression model, the percentage of each SVC’s reliable variance that is explained by the behavior PCs decays rapidly with SVC dimension, such that only 16 ± 8 SVCs (mean ± 95% CI, n=6 recordings with at least 131,072 neurons) show significantly more variance explained by behavior than temporally shuffled data (indicated in red, p<0.05, two-sided t-test). Each line indicates one mouse recording. **F.** Saturation of predictability of neural SVCs from behavior PC with increasing neuron number. Using the linear, instantaneous model, the first 32 neural SVCs initially become more predictable from behavior PCs as the neuron number increases, yet the variance explained by behavior saturates around 10,000 neurons (individual recordings in gray). The remaining SVCs above 32 (orange) are not predictable from behavior at any neuron number. **G.** Neural variance is concentrated in fewer neural SVCs during epochs of spontaneous behavior. The cumulative percent of neural variance explained is shown as a function of the number of neural SVCs for two scenarios: in blue, timepoints when the animal was idle and the total motion energy was below a given threshold (see STAR Methods for details); and in brown, timepoints when the animal was exhibiting epochs of spontaneous motor behavior. Shown is the mean ± standard error of n=6 recordings with at least 131,072 neurons, which was significantly higher during behaving epochs than idle periods, p<0.05, paired t-test. **H.** The lowest, behavior-related SVCs represent a much greater fraction of the neural variance during epochs of motor behavior. The ratio of the normalized fraction of variance during behaving versus idle epochs is shown for each SVC. Shown is the mean ± standard error of n=6 recordings with at least 131,072 neurons. This is consistent with our observation that a few behavior-related SVCs dominate the neural activity patterns. **I.** A comparison of the three different model types: linear, instantaneous; nonlinear (multilayer perceptron, MLP), instantaneous; and linear, multi-timepoint (see STAR Methods for details). Including nonlinearities and history both improve the prediction of the first neural 32 SVCs, but only by ∼10-15%. All traces indicate mean ± SEM from n=6 recordings with at least 131,072 neurons. **J.** The distribution of SVCs’ percent variance explained by behavior PCs for each of the three models. The neural SVCs for each recording are sorted by the percent of their reliable variance that is explained by behavior. Multi-timepoint models (gray line) predict many more SVCs than the instantaneous models (black and blue lines). The temporally shuffled data for the linear, multi-timepoint models are shown in red. The lines of corresponding color at the top of the plot indicate the number of neural SVCs that exhibit significantly higher predictability than the shuffled data (p<0.05, two-sided t-test). All traces indicate mean ± SEM of n=6 recordings with at least 131,072 neurons. **K.** Visualization of volumetric activity patterns and their predictions from behavior. (i) Instantaneous facial motion energy corresponding to the predicted neural activity, corresponding to the facial region highlighted in A. (ii-iv) Example coarse-grained volumetric activity maps exhibiting: (ii), the predictions from behavior (iii), and their difference (iv) are shown for two example timepoints. The coarse-grained volumetric activity maps are created by binning neurons into voxels of size 25 x 25 x 8 µm and averaging activity within each bin (see STAR Methods for details), providing a coarse map of activity across the volume. The predictions from behavior are found by predicting the neural SVCs utilizing a linear, multi-timepoint model and then transforming the predicted SVCs back into the full neural space. Timepoint t_1_ corresponds to a moment during a grooming epoch, whereas t_2_ corresponds to an idle period with little motion. Shown is an example 5 x 6 x 0.5 mm bi-hemisphere recording. Each image in (ii)-(iv) represents the 3D mean intensity projections of the volume; top left: xy projection, bottom: yz projection, right: xz projection. The scale bars in (ii) correspond to 1 mm laterally for the xy projection and 250 µm axially for the xz projection. The dynamics of these activities are shown in Video 3.

We first aimed to infer the behavior-related versus internal activity encoded by the neural SVC dynamics from our mesoscale recordings containing 134,628 to 970,546 neurons. To do so, we utilized linear and instantaneous reduced-rank regression to predict neural SVCs from the first 256 behavior PCs (see STAR Methods for details), an approach that allowed us to quantify what fraction of the reliable variance in each neural SVC was behavior-related (Figure 3C). Visual inspection of the predicted neural SVCs (Figure 3D) indicated that the lowest neural SVCs were highly predictable from behavior PCs, while higher neural SVCs did not appear as predictable. This observation was consistent across mice (Figure 3E), with only on the order of ten SVCs exhibiting significantly greater predictability from behavior than shuffled data (16 ± 8 SVCs, mean ± 95% CI, with significantly higher predictability than temporally shuffled data, p<0.05, two-sided t-test).

To further understand how our observed unbounded scaling of reliable neuronal dimensionality would differentially affect the inference of behavior-related versus internal components, we next asked how the fidelity of behavior-related encoding depended on the number of sampled neurons. We previously found that the reliability of many neural SVCs drops dramatically as a function of sampled neurons (Figure 2E), suggesting that any behavior-related SVCs would similarly require large populations of neurons for optimal prediction. We found that the percentage of reliable variance in the first 32 neural SVCs explainable from the behavior PCs initially increased with the number of randomly sampled neurons, but ultimately saturated at population sizes on the order of 10,000 neurons (Figure 3F, gray lines). This is consistent with our previous observation that for neuronal population sizes of 10,000 or more neurons almost all of these first 32 neural SVCs exhibited >75% reliable variance (Figure 2E and Figure S2C).

While the absolute percentage of reliable variance of the first 32 neural SVCs explained by behavior varied widely across recordings (16% to 60%), its dependence on neuron number and its saturation around 10,000 neurons was consistent (Figure 3F). In contrast to this observation, we found that the remaining higher reliable neural SVCs were not predictable from the behavior PCs irrespective of the number of recorded neurons (Figure 3F, orange lines), demonstrating the internal nature of these higher order components. These results demonstrate that while a significant fraction of observed cortical activity is correlated with instantaneous behavior, the majority of the reliable neural SVCs that can only be identified in large-scale, cellular-resolution recordings were not related to instantaneous behavior and thus are expected to be involved in other ongoing internal mental processes. We further investigated the relationship between the neural SVCs and spontaneous behavioral dynamics by dividing the neuronal timeseries into epochs corresponding to when the animal was engaged in motor behavior and those when the animal was idle. We indeed found that the observed high-dimensional neuronal geometry persisted even in idle epochs when the animal was not engaging in any motor behavior; however, the neural variance was more concentrated in the first 16 neural SVCs during motor behavior (Figure 3G), as can visualized by comparing the ratio of normalized neural variance in each SVC during motor behavior versus idle epochs (Figure 3H). Thus, we conclude that our previously identified unbounded scaling of reliable dimensionality as a function of neuron number (Figure 2F-G) represents primarily an increase in the number of internal encoded variables, as opposed to behavior-related variables with a moment-to-moment behavioral correspondence.

To corroborate our above conclusion, we further tested whether our inability to predict higher order SVCs from behavior was due to the simplicity of our linear and instantaneous reduced-rank regression model. Thus, we investigated whether various extensions to our model including utilizing a nonlinear network such as a multilayer perceptron (MLP) or inputting multiple behavioral timepoints would result in an increase in predictability of the higher neural SVCs from behavior. The multi-timepoint models were fit by inputting various numbers of additional behavior PC timepoints, binned in 1 second intervals to prevent over-fitting, and identifying the optimal number of timepoints around the instantaneous activity for each neural SVC independently (see STAR Methods for details). We found that while all such improvements to the model could increase the percentage of reliable neural SVC variance explained by the behavior PCs (Figure 3I), only the multi-timepoint models were able to predict significantly more additional neural SVCs on average (Figure 3J, 129 ± 54 SVCs, mean ± 95% CI) than temporally shuffled data (p<0.05, two-sided t-test; see also individual mice in Figure S3D-E).

We also found that on average predictability from behavior was optimal for a multi-timepoint window that stretched from 6 seconds before to 3 seconds after the corresponding instantaneous neural activity (Figure S3F-H). To confirm that these additional higher neural SVCs indeed encode an integration of the recent behavior, we showed that they could not be predicted by finding an optimal lag for each of the neural SVCs individually (Figure S3I), as opposed to the shared common lag utilized in the previous linear, instantaneous models.

Finally, we visualized the predictions of these neural SVCs from behavior by transforming the predicted neural SVCs back into the neural space and plotting their volumetric activity maps (Figure 3K and Video 3, see STAR Methods for details). Across animals we found agreement between the coarse-grained neuronal activity patterns and those predicted from the behavior PCs during epochs of spontaneous behavior, resulting in low prediction errors and consistent with our observation that 45 ±13% of the reliable variance in the first 32 neural SVCs is predictable from behavior (Figure 3K (ii-iv), top row t_1_). In contrast, the fidelity of the neuronal predictions was highly reduced for idle epochs with low motion energy, resulting in higher levels of prediction error (Figure 3K (ii-iv), bottom row t_2_). This observation corroborates the notion that a significant fraction of the variance across (mainly higher order) neural SVCs represents activity unrelated to the animal’s behavior and instead could represent neural activity related to internal or cognitive computations.

Taken together, our results confirm the global and robust encoding of spontaneous behaviors by a low-dimensional manifold as had been previously observed in smaller population sizes and low-resolution imaging modalities. We discovered that almost all behavior-related information was encoded relatively few (∼16) neural SVCs which accounted for only half of the total neural variance (54 ± 12% of total variance, mean ± 95% CI of n=12 recordings). However, this means that more than half of the observed reliable variance in these lower dimensions, as well as almost all of the observed reliable variance in the remaining higher order neural SVCs, are *not* explained by the animal’s instantaneous behavior (Figure 3E). This suggests their role in carrying information that is partially hidden and lacks an immediate behavioral readout such as internal states (Anderson, 2016; Flavell et al., 2022; Gordus et al., 2015; Marques et al., 2020; Norimoto et al., 2020; Sten et al., 2021; Sternson, 2020) or longer-timescale, integratory features, for example those involved in motor planning (Afshar et al., 2011; Churchland et al., 2010) or cognitive decision making (Lin et al., 2020; Mante et al., 2013; Shadlen and Kiani, 2013; Wei et al., 2019).

### Reliable neural SVCs exhibit a continuum of timescales and spatial distributions

While the latent factors identified via dimensionality reduction provide a more compact description of the diversity of cortical activity patterns and the features they encode, in reality neural computations are carried out by the spatiotemporal dynamics of populations of neurons distributed across brain regions. Thus, we next asked whether these latent neural SVC signals exhibited dynamics at characteristic spatiotemporal scales that could provide insight into the computations they would perform.

First, we asked if the neural SVC dynamics exhibited a characteristic timescale. To address this question and to quantify the dominant timescales, we fit an exponential decay to each SVC’s autocorrelation, thus defining its timescale τ. We found that the SVCs displayed timescales across a wide range, from minutes to hundreds of milliseconds (Figure 4A). The measured autocorrelation timescale decayed with neural SVC number in all recordings (Figure 4B). We found that the lower, behavior-related neural SVCs had the longest timescales, on the order of seconds to hundreds of seconds, whereas the higher neural SVCs operated at timescales of seconds or less, down to the minimal observable timescale in our data, which was limited by both the volume acquisition rate and kinetics of the calcium indicator. To investigate the dependence of the observed timescales on neuron number, we again performed random neuron subsampling analyses and found that as the number of sampled neurons increased and more neural SVCs became reliable, the number of SVCs with functional timescales greater than the sampling period also increased (Figure 4C). Thus, not only were these higher SVCs statistically reliable as defined previously, but they displayed a continuum of timescales from the limit of our detection up to the order of seconds, much greater than that expected from the experimental shot noise which is independent across frames. To control for this observation, we temporally shuffled the neural activity, which removed this diversity of functional timescales (Figure S4A), reducing the observed timescales to only the two second timescale defined by the shuffling procedure (see STAR Methods for details). The reliable activity patterns encoding these timescales could further be visualized by reconstructing the full neural activity from a low-rank approximation utilizing various numbers of neural SVCs (Video 4). We found that while the first 15 neural SVCs indeed could account for longer timescale patterns of coactivation across many neurons (Figure 4D, F, and H), many more finer timescales and patterns of neural activity were accounted for by the higher order neural SVCs (Figure 4E, G, and I).

**Figure 4.**
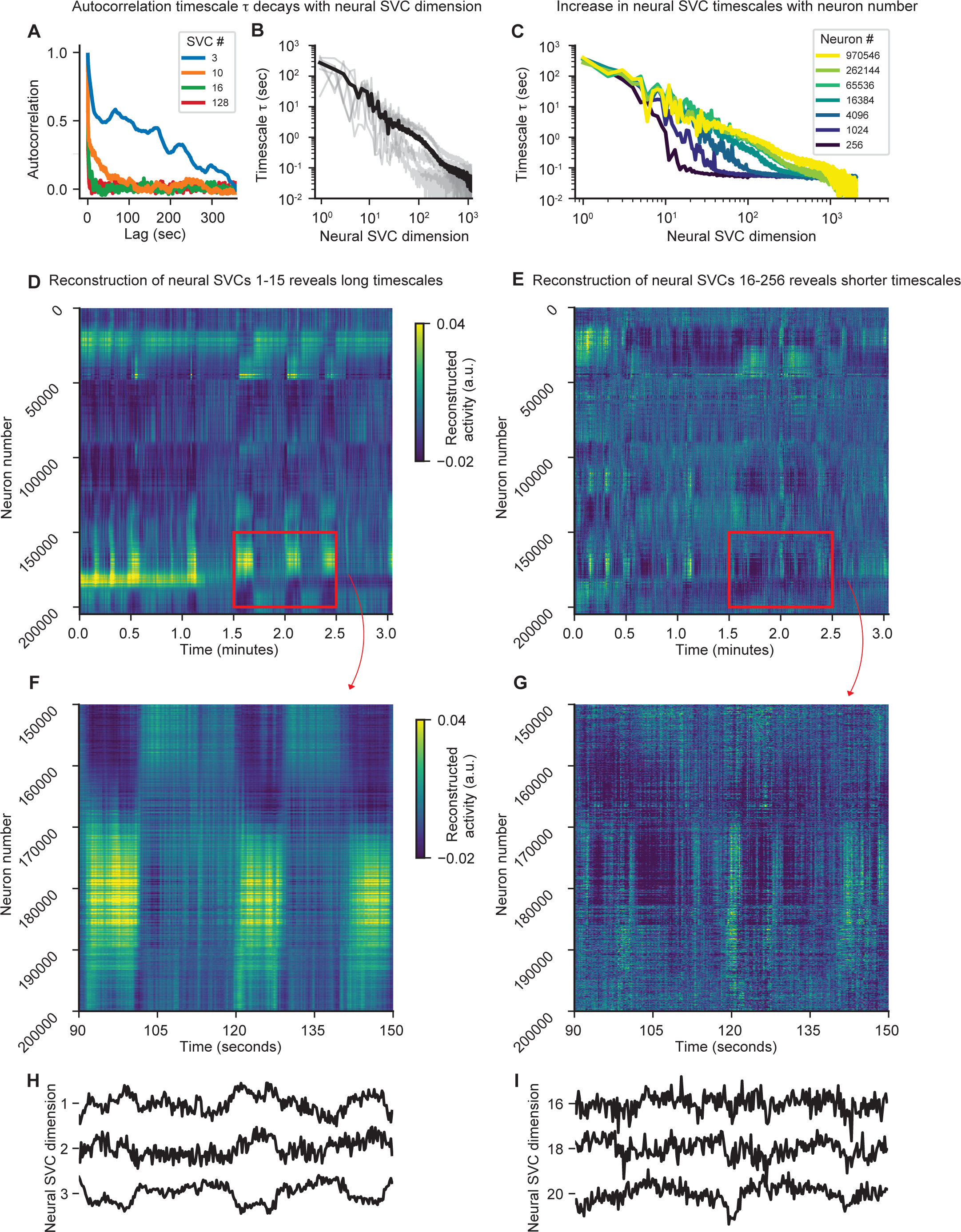
Latent neural SVC dynamics represent a continuum of timescales. **A.** Example autocorrelation curves of four neural SVCs from a recording containing 146,741 neurons. A wide variety of temporal scales are evident across all neural SVCs. **B.** Characteristic timescales within each neural SVC. The autocorrelation timescale τ, computed by fitting an exponential decay to each autocorrelation curve, decays with neural SVC dimension. Gray lines show n=6 recordings with at least 131,072 neurons, black indicates their mean. **C.** Characteristic timescales within each neural SVC increase as a function of the number of recorded neurons. As in B, but with a varying number of randomly sampled neurons. As the number of sampled neurons increases, more neural SVCs exhibit timescales greater than the period of our volume sampling rate. Shown as the mean timescales across n=6 full recordings with at least 131,072 neurons, with each neuron number randomly sampled four times per recording. **D - G.** Example heatmaps of reconstructed neural activity from neural SVC dimensions. The neurons are sorted utilizing the rastermap algorithm, which sorts neurons based on similarity of their temporal activity patterns. D. Reconstructed activity from neural SVCs 1-15. The information captured about the neural population within these SVCs visually exhibits longer timescale activity patterns. E. Reconstructed neural activity from SVCs 16-256. The contribution of these SVCs visually exhibits shorter timescale dynamics and a greater diversity of coactivation patterns across neurons. Shown are the reconstructions from the neural data displayed in Figure 1E. F and G display the red highlighted insets in D and E, respectively. **H** and **I.** Example neural SVCs representing distinct timescales. Corresponding timeseries for a few example neural SVCs used in the reconstructions in D and E, respectively.

Finally, we examined the spatial distribution of neuronal participation in each SVC by identifying the top fraction of neurons contributing to each SVC based on their normalized SVC coefficients (see STAR Methods for details). In an example cortical hemisphere recording, the top 3% of neurons participating in the lowest SVCs appeared hemisphere-wide yet spatially clustered, characterized by apparent clusters on the order of hundreds of microns distributed across many cortical regions (Figure 5A, (i) and (ii)). The spatial distribution of neurons contributing to the higher reliable neural SVCs, on the other hand, appeared much more uniformly distributed across the imaging volume (Figure 5A, (iii) and (iv); see also Figure S4B and Video 5). We quantified these spatial patterns by computing a local homogeneity index, which measured the average percent of neighbors within a given distance of a neuron that was also participating in that same neural SVC (see STAR Methods for details). The local homogeneity confirmed that the lower, behavior-related SVCs exhibited spatial clustering that decayed with distance, while the higher SVCs in some cases showed no homogeneity increase at small distances (Figure 5B). We also found a similar spatial distribution in our bi-hemispheric recordings (Figure 5C-D, Figure S4C and Video 6), which showed that nearly all of the neural SVCs were distributed in a cortex-wide fashion and were generally not limited to a single hemisphere or cortical region. We found that the local homogeneity was consistent across mice, with the first nearly 100 neural SVCs exhibiting some degree of local homogeneity, whereas temporally shuffled data exhibited no spatial structure (Figure 5E). Lastly, we confirmed that these higher SVCs were indeed widely distributed and not just encoded by a small, sparse group of neurons by computing the Gini index of the full neural weight vector for each SVC. The Gini index is a measure of sparsity or statistical dispersion that is widely used in econometrics, which was lowest and thus the least sparse for the higher SVCs, indicating their broad distribution across the entire imaging field (Figure S4D). Thus, together these spatial profiles made up a diverse repertoire of cortex-wide functional connectivity patterns which have been resolved at single neuron resolution for the first time. This combination of a continuum of timescales and a diversity of brain-wide spatial neural activity profiles likely represents a broadly-distributed functional network that underlies the transmission and manipulation of information throughout the cortex in order to enable the various internal computations and adaptive behaviors produced by the mammalian brain.

**Figure 5.**
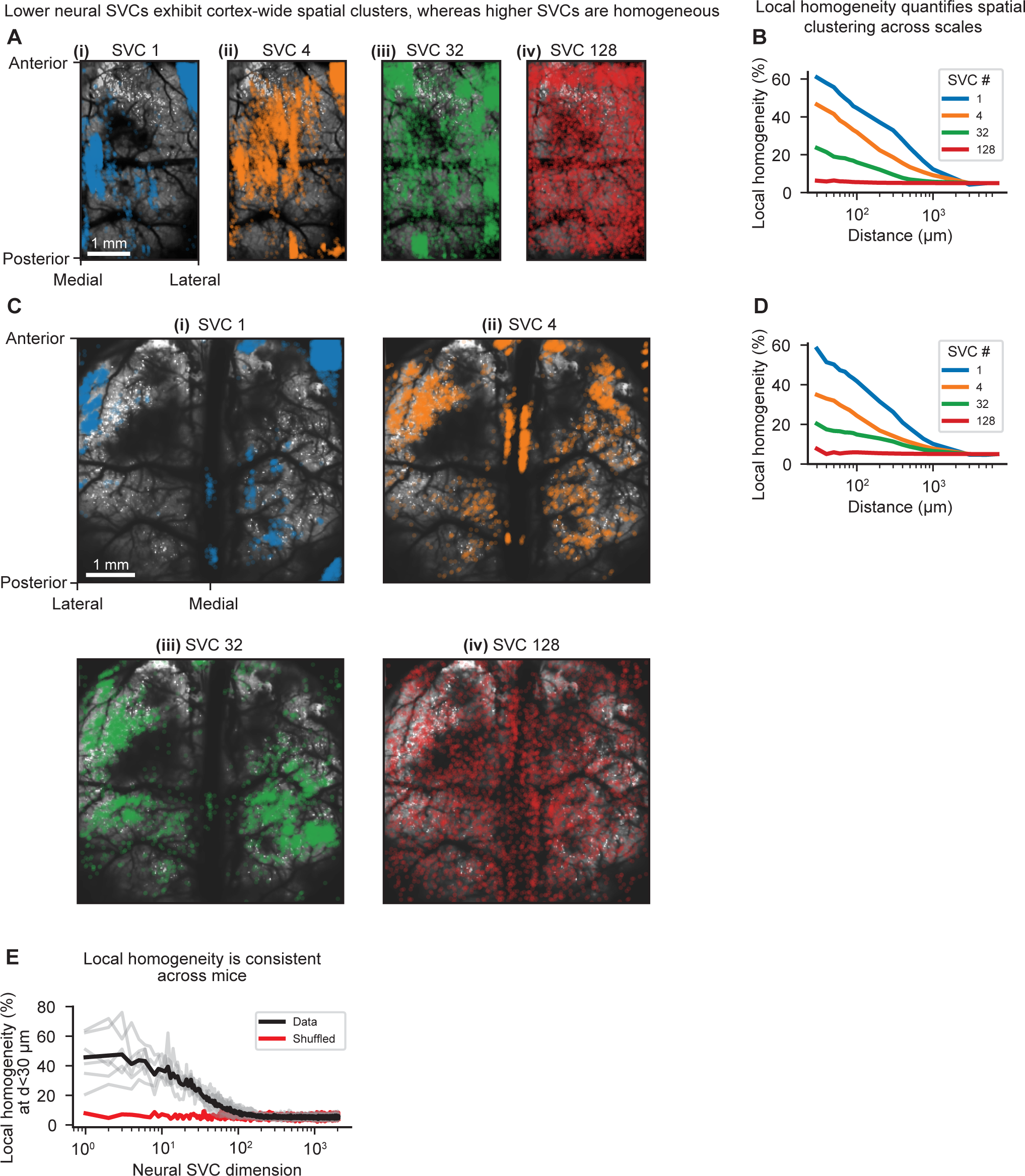
Lower and higher neural SVCs form distinct, spatially organized neuronal assemblies. **A.** Lower and higher SVCs exhibit distinct cortex-wide neuronal distribution profiles in a single hemisphere recording. The lateral spatial distribution of the top 3% of neurons contributing to four example neural SVCs in a 3 x 5 x 0.5 mm cortical hemisphere recording containing 315,363 neurons. Scale bar, 1 mm. A 3D rendering of the activity of the neurons contributing to each of these four SVCs is shown in Video 5. **B.** The local homogeneity, which quantities local spatial clustering of participating neurons, decreases with SVC number in a single hemisphere recording. The local homogeneity index, computed as the average percent of neighbors within a given distance that are also contributing to that same neural SVC, as a function of the radial distance for the four example SVCs in panel A. The lower neural SVCs exhibit strong local homogeneity at low distances, whereas the higher SVCs exhibit a flat profile with little to no clustering. **C.** Lower and higher SVCs exhibit distinct cortex-wide neuronal distribution profiles in a bi-hemispheric recording. The spatial distribution of the top 0.5% of neurons contributing to four example neural SVCs in a 5 x 6 x 0.5 mm bi-hemisphere recording with 970,546 neurons. The neurons contributing to each SVC are generally not restricted to single cortical regions or hemispheres, but show a wide distribution across both cortical hemispheres. Scale bar, 1 mm. A 3D rendering of the activity of the neurons contributing to each of these four SVCs is shown in Video 6. **D.** The local homogeneity, which quantities local spatial clustering of participating neurons, decreases with SVC number in a bi-hemispheric recording. The local homogeneity index as a function of distance shown for the bi-hemispheric example neural SVCs in panel C. **E.** The local homogeneity index profiles are consistent across mice. The local homogeneity computed at a radial distance of 30 µm for n=6 full recordings with at least 131,072 neurons. Each recording is shown in gray and their mean is showed in black. The mean of n=6 temporally shuffled recordings is shown in red.

## DISCUSSION

Despite technological advancements over the last decade that have pushed the number of simultaneously recorded neurons by several orders of magnitude across species, the emerging notion in an increasing body of literature is that the relevant dynamics of such large-scale neuronal dynamics are confined to low-dimensional manifolds that reflect neural computation, particularly the encoding of stimuli (Gervain and Geffen, 2018; Machado et al., 2022; Mante et al., 2013; Stopfer et al., 2003; Wu et al., 2018) and behavioral states (Dunn et al., 2016; Gallego et al., 2017; Kato et al., 2015; Sani et al., 2019). If true, on one hand this would beg the question of the biological utility of such a highly redundant and metabolically costly encoding scheme, and on the other hand would raise the question to what extent technological advances aimed at further increasing the number of recorded neurons would be necessary.

In this study, we hypothesized that in many cases the observed low-dimensional neural encoding is a reflection of technological limitations in the size of recorded population, ad hoc choices in the dimensionality reduction analysis, or a focus on only a subset of the population dynamics. Indeed, recent studies have utilized large-scale recording to highlight the multi-dimensional geometry of neural population dynamics (Dunn et al., 2016; Gallego et al., 2017; Kato et al., 2015; Sani et al., 2019; Schaffer et al., 2021), but there is a lack of data to address how the measured dimensionality depends on population size, recording modality, and a variety of other factors. Based on these hypotheses and using the unprecedented capability of our recently developed Light Beads Microscopy (LBM) to record up to one million neurons at multi-Hertz rate, we tested the conjecture that the measured neuronal dimensionality would continue to grow as a function of the size of sampled neuronal population.

To quantify this scaling, we utilized SVCA to identify dimensions of shared neural variance (the SVCs) and quantify the reliability of each dimension, thus providing an estimate of the planar neuronal dimensionality as the number of significantly reliable neural SVCs. Importantly, rather than the popular technique of choosing a dimensionality that represents some (usually) arbitrary fraction of the variance in the data, we perform a statistical significance test for each dimension independently, thus defining the reliable dimensionality. We demonstrate for the first time that the reliable dimensionality in our recordings scales in an unbounded fashion and according to a power law with the number of neurons. This observation indicates that even further gains in neuron count and imaging FOV should uncover additional latent components. In fact, our results suggest that we are still experimentally limited by nearly every relevant imaging parameter, such that increased dimensionality would also be expected with further increases in recording duration and speed, even though the relative gains in dimensionality across these parameters would be different. Although saturation in dimensionality is ultimately expected at the whole-brain scale, current cellular-resolution technologies are still far from whole-brain rodent imaging and will require highly scalable recording modalities that can penetrate thick tissue with minimal invasiveness.

Next, by establishing the relationship between the reliable neural SVCs in our recordings and the animal’s epochs of spontaneous and uninstructed behavior, we identified a low-dimensional encoding of spontaneous behavior within 16 ± 8 of the largest neural dimensions. While this observation is consistent with recent studies that have shown a broad representation of motor behavior across the cortex and brain (Kaplan and Zimmer, 2020), our work extends these in important ways. First, while previous studies have speculated that motor behavior is encoded by a low-dimensional subset of the latent cortical dynamics, our systematic study of scaling, combined with a 1000-fold increase in neuron count relative to the previous largest experiments (Weisenburger et al., 2019), provides strong evidence that this low number of behavior-related dimensions is a cortex-wide property and not a limitation of the experimentally feasible population size or neural volume sizes and rates. Second, our large-scale, simultaneous recording technology enabled us to identify the cortex-wide cellular ensembles contributing to each neural dimension. We discovered that each behavior-related dimension was encoded by multiple spatial clusters of neurons distributed across cortex, likely representing a structured brain-wide network broadcasting the behavioral state of the animal at any given time. Finally, in addition to such instantaneous correlations with behavior, we found that more than 100 additional neural dimensions can be predicted above chance from a longer ten second multi-timepoint window that includes a history of behavioral activity, highlighting the importance of considering the longer timescale dynamics of spontaneous behavior and neural activity. We speculate that these additional dimensions reflect neural computations that are temporally structured at timepoints near movement, but not instantaneously correlated with them, such as motor planning and decision making (Briggman et al., 2005; Inagaki et al., 2018; Svoboda and Li, 2018).

Most importantly, our observation of a low-dimensional subspace encoding behavior-related information demonstrates that the largest gains from large-scale recordings come from the ability to robustly infer purely internal neural dynamics. We found that >90% of the reliable neural dimensions showed no temporal correspondence with the epochs of spontaneous motor behavior. We thus hypothesize that they encode purely internal variables without a consistent, time-locked behavioral readout, or potentially spontaneous activation (Luczak et al., 2009; Pezzulo et al., 2021; Romano et al., 2015) of sensory and other non-motor cell ensembles. In addition to their statistical reliability, these higher order internal SVCs exhibited fast-timescale, spatially homogeneous dynamics, which we show are distinct from noise and may underly a brain-wide network conveying internal variables. This apparently homogeneous spatial distribution suggests the higher neural SVCs may act via a brain-wide circuitry or large-scale neuropeptide modulation. Further, this diversity of cortex-wide spatiotemporal patterns may enable flexible encoding and computations across the entire range of circuit motifs required for generating adaptive behavior.

However, we show that in order to reliably extract these high-dimensional features, population sizes of tens to hundreds of thousands of simultaneously recorded neurons are required. Considering that simultaneously recorded neuronal population sizes of such scale have only very recently become experimentally attainable, only very few comparable experiments are available that provide context for these signals in other tasks or behaviors. Additionally, it is still debated how spontaneous activity is related to evoked activity or neural dynamics underlying goal-directed behavior (Avitan et al., 2021; Sussillo and Abbott, 2009). We expect further studies using large-scale simultaneous neurorecording technologies will reveal whether these spontaneous spatiotemporal dynamics form a set of generalized cortical computations that are re-used across tasks, as previously argued using low-resolution widefield recordings (MacDowell and Buschman, 2020).

Thus, while our LBM data add to the growing evidence that the encoding of behavior occurs in a widely broadcasted manner that may facilitate the integration of sensory inputs with motor variables as early as primary sensory cortex, we speculate that our observed higher SVCs may underlie a similar brain-wide network which broadcasts purely internal variables, such as attention, motivation, hunger or thirst, and fear. These variables often primarily serve to modulate the ways in which neural circuits, and thus the animal’s behavior, respond in a dynamic and adaptive fashion to an ever-changing sensory environment, and thus may need to be combined with sensory, motor, and other signals that are encoded and manipulated within circuits distributed across the entire brain. We expect that the combination of these large-scale recording technologies with ethologically relevant tasks and data-driven quantification of behavior provide a promising avenue to understand how these internal variables are combined with sensory and motor representations to shape behavior across trials and over a lifetime.

Finally, we note that the techniques implemented here provide a data-driven estimate of the planar dimensionality, and a more general estimation of the intrinsic dimensionality of complex systems has proven challenging (Altan et al., 2021; Bialek, 2022; Ganguli and Sompolinsky, 2012). For instance, the relationship between latent dimensionality and neural population size is expected to also be a function of the complexity of the animal’s environment and tasks (Jazayeri and Afraz, 2017), and in our study we have focused on spontaneous dynamics while the animals were isolated and in the dark. Additionally, although by far the most common approach, our focus on the planar dimensionality does not account for the possibility that the intrinsic dimensionality of cortical dynamics may be lower if it should be found to lie on a highly nonlinear manifold. However, nonlinear manifold inference is generally computationally challenging, especially on high-dimensional, noisy data, and often requires assuming *a priori* a simple topological structure (Chung and Abbott, 2021; Cunningham and Yu, 2014; Gardner et al., 2022). Nonetheless, the planar dimensionality, also referred to as embedding dimensionality, has been argued to be important for how information is processed by neural circuits (Jazayeri and Ostojic, 2021), particularly given the wealth of evidence suggesting that sensory and motor systems, and recently also more abstract cognitive representations, actively create representations that utilize orthogonal subspaces and linear decodability (Bernardi et al., 2020; Chung and Abbott, 2021; DiCarlo and Cox, 2007). We expect that further mathematical and computational advancements will be made towards a more systematic inference of nonlinear manifolds and that the development and validation of these tools would benefit from our large-scale recording capabilities.

## ACKNOWLEDGEMENTS

We thank Q. Lin and S. Lu for comments on the initial versions of the manuscript, the members of the Vaziri lab for discussions regarding data management and analysis, and the Rockefeller University High Performance Computing Cluster for compute access. Research reported in this publication was supported by the National Institute of Neurological Disorders and Stroke of the National Institutes of Health under award numbers 1U01NS115530, 5U01NS103488 and 1U01NS126057, and the Kavli Foundation through the Kavli Neural Systems Institute.

## AUTHOR CONTRIBUTIONS

J.M. contributed to the project conceptualization, performed experiments, analyzed data, and wrote the manuscript. J.D. contributed to the behavioral recording setup and performed imaging experiments. H.K. and F.M.T. performed cranial window surgeries. A.V. conceived and led the project, designed the experiments and data acquisition approach, guided data analysis, and wrote the manuscript.

## SUPPLEMENTARY FIGURES

**SI Figure related to Figure 1. Example recording configurations used in this study.**

**A - C.** Example data from various recording configurations, displaying the following: (i): Top: Imaging FOV denoted on the Allen brain atlas. Bottom: Standard deviation intensity projection from an example plane. (ii): Neuronal heatmap for 3 minutes from the recording, sorted using rastermap. (iii): Fifteen example neuronal timeseries from the red highlighted region in (ii). (iv) and (iv): Simultaneous timeseries of each of the first three behavior PCs, aligned to the neuronal activity in (ii) and (iii), respectively. A. Example data from a 1.2 x 1.2 x 0.5 mm recording at 9.6 Hz and containing 21,087 neurons. B. 3 x 5 x 0.5 mm single hemisphere recording at 4.7 Hz and containing 204,798 neurons. C. 5.4 x 6 x 0.5 mm bi-hemispheric recording at 2.2 Hz and containing 970,546 neurons.

**D.** 3 x 5 x 0.5 mm single hemisphere recording at 4.7 Hz and containing 174,113 neurons. (i): Full neuronal timeseries heatmap for an example 30 minutes of recording. (ii) and (iii): subsequent zoomed insets, as denoted by the red highlighted regions.

**SI Figure 2, related to Figure 2. Additional neural SVCA results.**

**A.** The reliability of many SVCs increases with the number of recorded neurons. The reliability of each neuronal SVC can be visualized by comparing the two neural sets’ SVC projections. Shown are example z-scored SVC projections from both neural sets from populations with various numbers of neurons, from 2,048 to 131,072 neurons. The percentage of reliable variance over the full testing timepoints is shown to the left of each trace.

**B.** Temporally shuffled data do not contain reliable SVCs. As in panel A, except utilizing the temporally shuffled neuron timeseries (see STAR Methods for details).

**C.** Scaling of reliable dimensionality for different reliable variance thresholds. The scaling of reliable dimensionality with neuron number is consistent across a range of arbitrary thresholds of percent reliable variance. The number of SVCs greater than this minimum percentage of reliable variance is shown as the mean ± standard deviation of n=12 recordings.

**D.** Normalized reliable dimensionality as a function of volume acquisition rate. Reliable dimensionality (normalized for each recording to a maximum of 1), measured after post-hoc temporal downsampling, increases with imaging framerate for our n=3 fastest recordings at 10 Hz and 1.2 x 1.2 x 0.5 mm FOV. Temporal downsampling is performed by removing the i-th frame from each recording, with i varying from 2 to 14.

**E.** The reliable dimensionality as a function of neuron number and the duration of the recording. The dimensionality does not strongly depend on the recording duration except in the case of very large neuron number (light green and yellow lines). Shown as the mean ± standard deviation of the number of reliable SVCs for n=6 recordings with at least 131,072 neurons, with 10 neuron samplings per recording.

**F.** Reliable variance spectra exhibit a continuous power law decay. Various numbers of neurons are subsampled randomly from across the entire imaging FOV of a total recording of one million simultaneously recorded neurons (Fig 2E). The total reliable variance spectrum exhibits a power law decay with no significant changes of the variance spectra as a function of neuron number. The reliable variance shown is the mean over n=10 random samplings and normalized to sum to 1.

**G.** Variance-based dimensionality estimates, such as PCA, do not exhibit unbounded scaling. Utilizing the traditional PCA-based approach without cross-validation to measure reliability, the number of PCs required to represent a given percentage of the total variance is plotted versus neuron number, indicating a clear bounded scaling in the case of all three thresholds. The color indicates the required percentage of variance explained and each line is a separate mouse recording. Shown is the mean number of PCs over n=10 random samplings.

**H.** The scaling of the reliable dimensionality does not depend on the density of neuronal sampling. Each volume size is randomly placed within the full FOV and corresponds to a square of given size laterally and the full 500 µm axial range. The reliable dimensionality is shown as the mean ± SEM over all n=12 recordings that had a sufficiently large volume.

**SI Figure 3, related to Figure 3. Quantification of behavior and prediction of neural SVCs from multi-timepoint behavior.**

**A.** The behavior PCs represent a multi-dimensional description of behavior. The cumulative percent variance explained as a function of the number of behavior PCs from n=12 recordings. This behavioral decomposition is performed using Facemap, which performs PCA on the motion energy between consecutive frames. The behavior appears multi-dimensional and relatively consistent across mice.

**B.** A number of clustered behavioral motifs are evident in the behavior PC dynamics. A t-SNE embedding, shown as a probability density function (PDF), of behavior PC profiles for an example recording. This behavioral map contains a number of dense clusters, denoting commonly performed movements, which correspond to unique motion energy profiles that are reconstructed in panel B.

**C.** Example clustered behavioral motifs. The average motion energy pattern reconstructed from the mean behavior PC profile of each cluster shown in panel B indicating association with specific behavioral motifs. The color of the bounding box corresponds to the region denoted in panel B.

**D** and **E.** The percent of reliable variance that is explained by behavior for each neural SVC for single time-point (D) and multi-timepoint (E) models across n=12 recordings. Measurements are shown in black and the temporally shuffled data is shown in red. Panel D contains the predictions from instantaneous behavior, while panel E contains the predictions from an optimal multi-timepoint behavior window (calculated over 11 seconds before to 6 seconds after the instantaneous behavior, see STAR Methods for details) identified for each SVC independently.

**F-H.** The optimal multi-timepoint behavioral window for neural prediction is about 9 seconds long. In F, the average percent variance explained by behavior for the first 50 SVCs (across n=6 recordings with at least 131,072 neurons) is shown as a function of the number of input behavioral timepoints, with a given number of seconds in the past and future relative to the instantaneous lag. The optimal window is circled in black, which occurs from 6 seconds before to 3 seconds after the instantaneous behavior. G and H show the variance explained as a function of the number of seconds before and after neural activity, respectively.

**I.** An optimized instantaneous lag for each neural SVC does not improve the model performance. The mean ± SEM percent variance explained across n=6 recordings with at least 131,072 neurons is shown for the linear, instantaneous model with a common shared lag (black), the linear, multi-timepoint model (gray), and a lag-optimized linear, instantaneous model (orange), where a single lag between the behavior PCs and neural SVC is optimized for each SVC independently (see STAR Methods for details).

**SI Figure 4, related to Figures 4 & 5. Additional neural SVC spatiotemporal characteristics.**

**A.** Reliable neural SVCs exhibit distinct timescales from temporally shuffled data. The neural SVCs (black) exhibit a continuum of timescales. The temporally shuffled data (red) exhibit timescales primarily around two seconds, which is the interval at which the neural data is chunked during the shuffling procedure (see STAR Methods for details). Shown are the means for n=6 full recordings with at least 131,072 neurons.

**B.** Lower and higher SVCs exhibit distinct cortex-wide neuronal distribution profiles in a single hemisphere recording. The lateral spatial distribution of the top 3% of neurons contributing to four example neural SVCs in another 3 x 5 x 0.5 mm cortical hemisphere recording containing 146,741 neurons. Scale bar, 1 mm.

**C.** Lower and higher SVCs exhibit distinct cortex-wide neuronal distribution profiles in a bi-hemispheric recording. The spatial distribution of the top 0.5% of neurons contributing to four example neural SVCs in another 5 x 6 x 0.5 mm bi-hemisphere recording with 703,439 neurons. Scale bar, 1 mm.

**D.** The higher order neural SVCs exhibit the least neuronal sparsity, indicating they are encoded by large populations of cortex-wide neurons. The Gini index (a measure of sparsity in which 0 indicates complete uniformity and 1 indicates maximal sparsity) of the vector of SVC coefficients as a function of SVC number. Sparsity peaks in the first 10s of SVCs, while the higher order SVCs exhibit the lowest sparsity. Each recording is shown in gray and their mean is showed in black for n=6 full recordings with at least 131,072 neurons.

## STAR METHODS

## RESOURCE AVAILABILITY

### Lead contact

Further information and requests for resources should be directed to the lead contact, Alipasha Vaziri.

### Materials availability

This study did not generate new unique reagents.

### Data and code availability

All of the code and processed data used to perform analyses and generate figures will be made available in a public repository upon acceptance of the manuscript. All other information required to reanalyze the data reported in this paper is available from the lead contact upon reasonable request upon acceptance of the manuscript.

## EXPERIMENTAL MODEL AND SUBJECT DETAILS

### Animal subjects and surgical procedures

All surgical and experimental procedures were approved by the Institutional Animal Care and Use Committee of The Rockefeller University. VGlut1-cre x fl-GCaMP6s or fl-GCaMP6f (Jackson Labs stock numbers 023527, 031562 and 030328, respectively (Daigle et al., 2018)) crossed mice were bred in house. Cranial window implantation was performed as previously described (Demas et al., 2021), during which a circular 8mm diameter dual-hemisphere craniotomy was performed. The cranial window was formed by implanting a circular 8 mm glass coverslip with 1 mm of the bottom removed (#1 thickness, Warner instruments) before sealing with tissue adhesive (Vetbond). Mice with significant regrowth or otherwise unclear windows were euthanized and not used for imaging experiments. All mice were 30-95 days of age at the time of first procedure and were 47-170 days old during imaging experiments. Mice were allowed food and water *ad libitium*.

## METHODS DETAILS

### Imaging parameters

Data acquisition was performed utilizing LBM as described by (Demas et al., 2021), utilizing a custom multiplexing module interfaced with a commercial mesoscope (Thorlabs, Multiphoton Mesoscope). n=6 total mice, male or female, were imaged across n=12 sessions for a duration of either 30 minutes (n=2) or 60 minutes (n=10). Three FOV types were recorded: 1.2 x 1.2 x 0.5 mm, 2 µm lateral pixel spacing, at 10 Hz volume rate (yielding 6,519 – 21,087 neurons); 3 x 5 x 0.5 mm, 5 µm spacing, at 4.7 Hz (134,628 – 315,363 neurons); and 5.4 x 6 x 0.5 mm, 5 µm spacing, at 2.2 Hz (703,439 – 970,546 neurons). Imaging power was restricted to <250 mW in the smallest FOV (1.2 x 1.2 x 0.5 mm) while larger recordings ranged from 289 – 415 mW. These power settings remained within previously demonstrated safe thresholds for heat induced immunohistochemical reactions and were shown to be insufficient to drive astrocyte activation marker GFAP immunoreactivity (Demas et al., 2021).

Mice were head-fixed on a low-friction, self-driven belt treadmill (Jackson et al., 2018) with a rotation encoder affixed to the rear axle (Broadcom, HEDR-5420-ES214) to measure tread position during recordings. Treadmill position, the microcontroller clock value, and blinking status of an infrared LED were streamed to the control computer via a serial port connection and logged with a separate data logging script. The treadmill control software is available at https://github.com/vazirilab/Treadmill_control.

### Facial videography

Infrared LEDs (850 nm) illuminated the mice to enable recording of their spontaneous movements while head-fixed yet able to freely move on the treadmill. Videos were acquired at 30+ Hz using a camera (FLIR BFS-U3-51S5M-C Blackfly S) pointed at one side of the mouse’s face and body, equipped with a zoom lens (Navitar MVL7000) and infrared filter (Midwest Optical BN850; 850nm with 45nm FWHM) to reject two-photon laser excitation and other stray light. Behavior PCs quantifying the animal’s behavior were calculated utilizing Facemap (Stringer et al., 2019b) version 0.2.0 on a region of interest centered on the face of the mouse (for example, see Figure 3A-B) with a spatial binning of two pixels. The behavior PCs represented the PCA decomposition of the motion energy, defined as the absolute pixel-wise difference between each consecutive camera frame. In Figure 1G-H, the total motion energy is calculated as the sum of the absolute pixel-wise difference between consecutive frames. Behavior data were synchronized with neural recordings by monitoring an LED blinking every five seconds controlled by the master data acquisition software. Behavior PCs were then re-sampled to match the neural timestamps using linear interpolation.

## QUANTIFICATION AND STATISTICAL ANALYSIS

### Data processing

Raw calcium imaging data were processed as described previously (Demas et al., 2021) utilizing a custom pipeline based on the non-rigid version of NoRMCorre motion correction (Pnevmatikakis and Giovannucci, 2017) and the patched, planar version of the CaImAn software package (Giovannucci et al., 2019; Pnevmatikakis et al., 2016). The full pipeline is available at https://github.com/vazirilab/MAxiMuM_processing_tools. The spatial correlation threshold was held at the default value of 0.4 and the minimum signal-to-noise parameter was set to 2. In contrast to previous applications of SVCA (Stringer et al., 2019b), we do not bin our neural time series past the native imaging framerate, and all subsequent analyses are performed at the framerate specified for each FOV type.

### Shared variance component analysis (SVCA)

SVCA was performed as previously described (Stringer et al., 2019b) utilizing a custom Python package. The SVCA algorithm is based on applying maximum covariance analysis to two subsets of the neuronal population, with cross-validation in time. Here, we sketch the SVCA procedure, which provides a lower-bound estimate for the proportion of variance in a neural population that is reliably encoded by low-dimensional latent dynamics. The full population was in some cases subsampled to a given neuron number *N* by either: 1. randomly selecting neurons across the full FOV (Figures 2D-F, S2A-C, S2E-F, 3F, 3I, 4C); 2. sampling an expanding FOV comprised of the *N* neurons closest to the center of the imaging volume (Figure 2G); or 3. sampling *N* neurons from a lateral FOV of a given size randomly placed within the FOV (Figure S2H).

The selected population is split into two sets by dividing the imaging FOV laterally into squares of 250 µm and collating non-adjacent squares in a checkerboard pattern into one set, to prevent any axial crosstalk between sets due to out-of-focus fluorescence. Each neuron’s activity was z-scored. Timepoints were separated by chunking the data into 72 second intervals and randomly assigning each to either the training or testing set. In any case where the number of neurons *N* in a set was greater than the number of training timepoints *T*, PCA was applied on each set’s data matrix and all *T* PCs were kept to reduce the dataset to the rank of the matrix and decrease memory requirements; since all PCs were kept, this is mathematically equivalent to performing the following analysis on the full matrices, but more memory efficient. The covariance matrix between the two neuron sets c= F_train_ G_train_^T^/T-1 was decomposed using a randomized singular value decomposition (SVD) estimator (Pedregosa et al., 2012) to yield 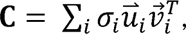, where 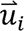 and 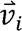 correspond to the *i*-th maximally covarying projections of each neural set. The reliable variance in the *k*-th SVC is then given by

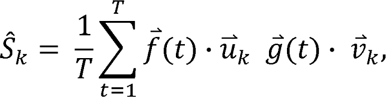

where 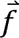 and 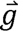 are the held-out testing timepoints. This is then normalized by the arithmetic mean of the total variance s_k,tot_ in both neural sets to calculate the percentage of reliable variance in each component.

To estimate the reliable dimensionality, we consider the *i*-th SVC to be significantly reliable if its percentage of reliable variance is greater than four standard deviations above the mean of the *i*-th SVC of time-shuffled datasets. This in all cases turned out to be stricter than performing a two-sided t-test requiring p<0.05. Temporal shuffling was performed by chunking each neuron’s timepoints into chunks of two seconds in duration and then randomly permuting these chunks, removing temporal alignment between neurons. This two second chunking and shuffling was chosen so as to sufficiently remove all temporal alignment among neurons while preserving some of the statistics of the calcium kernel. This was consistent with shuffling performed by random circular permutation of each neuron’s timeseries independently when the number of neurons was less than the number of timepoints; however, these circular permutation shuffling approaches can no longer remove all alignment when the number of neurons is significantly greater than the number of timepoints.

Post-hoc temporal down-sampling analyses (Figure S2D) were performed by removing every *i*-th timepoint (from *i*=1 to 14) and repeating the SVCA procedure. Post-hoc duration analyses (Figure S2E) were performing by keeping only the first *D* minutes of the original one hour recordings.

In Figure 3G-H, the total variance was calculated as described above but only including timepoints from the test set where the animal was either idle or exhibiting an epoch of spontaneous behavior. Idle and behaving epochs were separated by thresholding the total motion energy at a chosen level that visually separated epochs of movement versus non-movement, and any idle timepoints within 1 second of a movement epoch were ignored. The variance in each epoch was normalized to sum to 1, representing the fraction of variance in each neural SVC, before computing the cumulative fraction of variance (Figure 3G) or the ratio between the variance in behaving versus idle epochs (Figure 3H) for each SVC.

### Predicting neural SVC activity from behavior PCs

To estimate the amount of behavioral information contained with the SVCs, we utilized reduced-rank linear regression to predict the SVCs (F_train_ and G_train_) from the first 256 mean-subtracted behavior PCs (x_train_), as described previously (Stringer et al., 2019b) utilizing a custom Python package. The behavior PCs were shifted by a lag of approximately 200 ms (0 frames for 2.2 Hz recordings, 1 frame for 4.7 Hz, and 2 frames for 10 Hz) to account for the rise time of the calcium signal. The residual error of each SVC

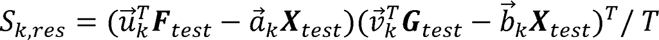

quantifies the amount of reliable variance in the *k*-th SVC that cannot be predicted from the behavior such that 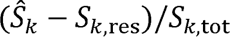 represents the fraction of reliable variance explainable by the behavior PCs. The rank was varied from 1 to 64 and the rank with optimal variance explained was displayed.

Additional models were tested similarly. Nonlinear regression was performed with multilayer perceptrons containing a single hidden layer with 1 to 64 units, ReLU activation, and trained using the Adam optimizer (Kingma and Ba, 2014; Pedregosa et al., 2012). For multi-timepoint models, additional inputs were provided to the model by averaging behavior PC activity in 1 second bins from up to 11 seconds before to 6 seconds after the instantaneous neural activity.

This 1 second binning was utilized in order to prevent model overfitting. The optimal model was found for each neural SVC independently after varying the number of time bins provided as input. Shuffling tests for prediction from behavior were performed by predicting the shuffled neural SVCs (see above) from the original behavior PC activities.

### Data visualization

Raw two-photon calcium images were denoised using DeepInterpolation (Lecoq et al., 2021) to improve the visualization of neurons in Video 2 and the standard deviation projection images throughout the manuscript. A unique DeepInterpolation model was trained until convergence for each recording, utilizing the “unet_single_1024” network configuration and with the number of pre and post frames set to roughly 1 second. Calcium traces were individually normalized via z-scoring to improve visualization. Line plots of individual calcium traces were smoothed with a one-second moving average to improve transient visualization. Heatmaps of calcium dynamics were sorted using the Python version of the rastermap algorithm (Stringer et al., 2019b). The authors’ analyses and visualizations were built using the open source Python scientific computing ecosystem, including Matplotlib (Hunter, 2007), NumPy (Harris et al., 2020), SciPy (Virtanen et al., 2020), and Scikit-learn (Pedregosa et al., 2012); we are indebted to their many contributors, maintainers, and funders.

The coarse volumetric activity maps in Figure 3K and Video 3 were visualized by binning the neurons into 25 x 25 x 8 µm coarse voxels. The activity of the neurons within each bin is averaged at each timepoint, providing a coarse overview of the activity patterns across the full FOV. The maps were convolved with a gaussian filter with a standard deviation of 50 µm laterally and 16 µm axially to improve visualization and then z-scored.

The 3D renderings in Videos 5 and 6 were created by plotting each neuron as a sphere and visualizing its z-scored activity by the opacity, which was maximally opaque at a z-score of 5 or above and not shown at a z-score of −0.2 or below. The top 3% or 0.5%, for each video respectively, of the neurons contributing to the four example SVCs were shown.

## KEY RESOURCES TABLE

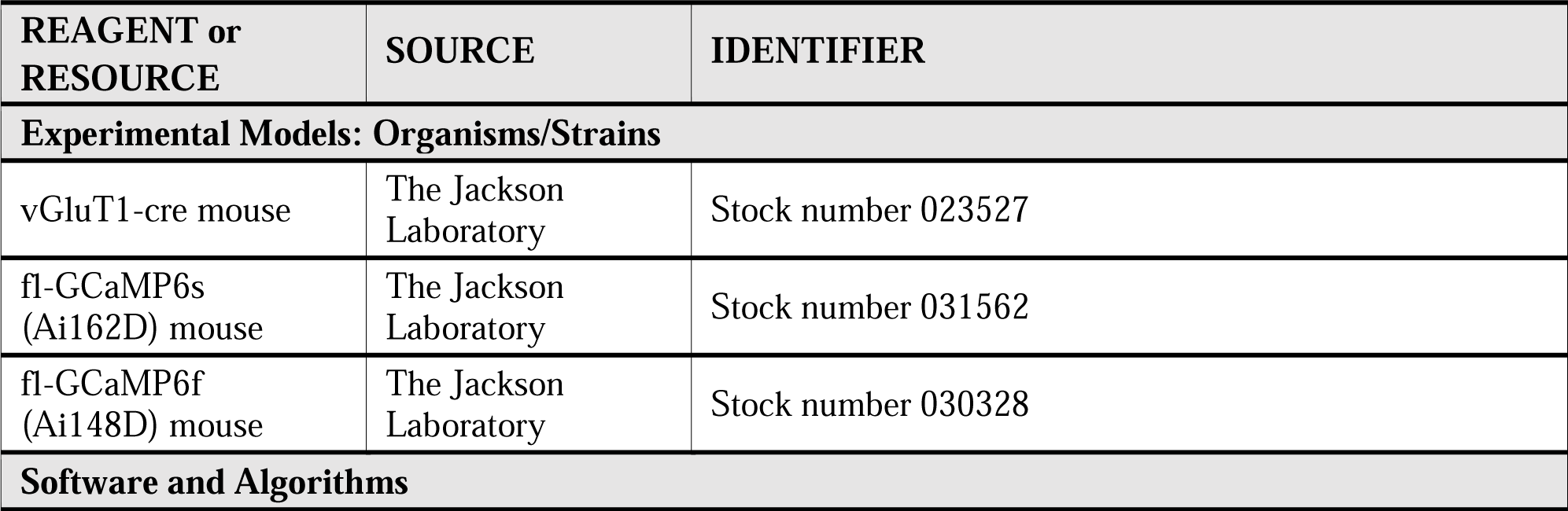
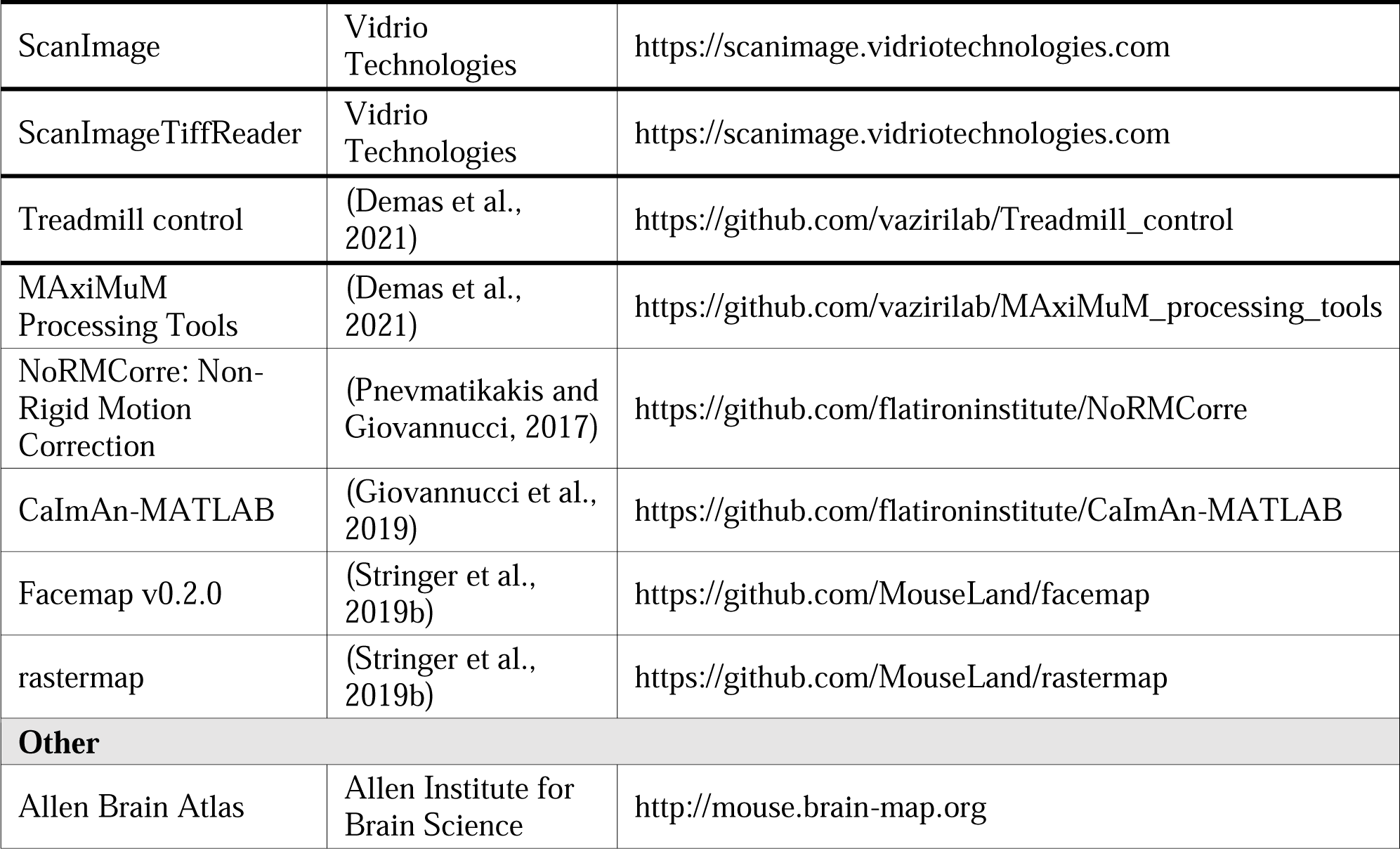

## SUPPLEMENTARY VIDEO TITLES AND LEGENDS

**Video 1. Example recording of a single plane (330 µm depth) in a 3 x 5 x 0.5 mm right cortical hemisphere recorded at 4.7 Hz.** This recording corresponds to a FOV similar to that in Figure 1D, covering parts of somatosensory, retrosplenial, posterior parietal, and visual cortices.

Playback sped up 5 times. Scale bar, 250 µm.

**Video 2. Example mouse behavior during a 1 hour imaging experiment.** The mice exhibited spontaneous epochs of whisking, grooming, running, and other behaviors.

**Video 3. Example volumetric activity maps and predictions from multi-timepoint behavior for the dataset shown in Figure 3I**. The volumetric activity maps are displayed as described in Figure 3I and STAR Methods, corresponding to the bi-hemispheric FOV indicated in Figure S1C(i). Left: the activity map from the raw neural data. The xy projection scale bar indicates 1 mm laterally and the xz projection scalebar indicates 250 µm axially. Middle: the predicted neural activity from behavior using a linear, multi-timepoint model. Right: the instantaneous motion energy, reconstructed from the first 500 behavior PCs. Playback sped up 5 times.

**Video 4. Example heatmap of reconstructed neural activity as a function of the number of SVCs utilized.** Shown is the reconstructed activity from the number of SVCs denoted above for the dataset shown in Figure 4D-E.

**Video 5. 3D rendering of 17,572 neurons representing the top 3% contributing to four example SVCs from the data shown in Figure 5A-B**. This recording corresponds to the single hemisphere FOV indicated in Figure 1D. Colors correspond to those in Figure 5 (SVC 1: blue, 4: orange, 32: green, 128: red). Each dot corresponds to one neuron, and its activity is represented by its opacity (see STAR Methods for details). Playback sped up 4 times.

**Video 6. 3D rendering of 19,408 neurons representing the top 0.5% contributing to four example SVCs from the data shown in Figure 5C-D**. This recording corresponds to the bi-hemispheric FOV indicated in Figure S1C(i). Colors correspond to those in Figure 5 (SVC 1: blue, 4: orange, 32: green, 128: red). Each dot corresponds to one neuron, and its activity is represented by its opacity (see STAR Methods for details). Playback sped up 10 times.

