## Supplemental Figure 1 for "Simultaneous, cortex-wide and cellular-resolution neuronal population dynamics reveal an unbounded scaling of dimensionality with neuron number"

**Figure S1**: related to Figure 1. Example recording configurations used in this study

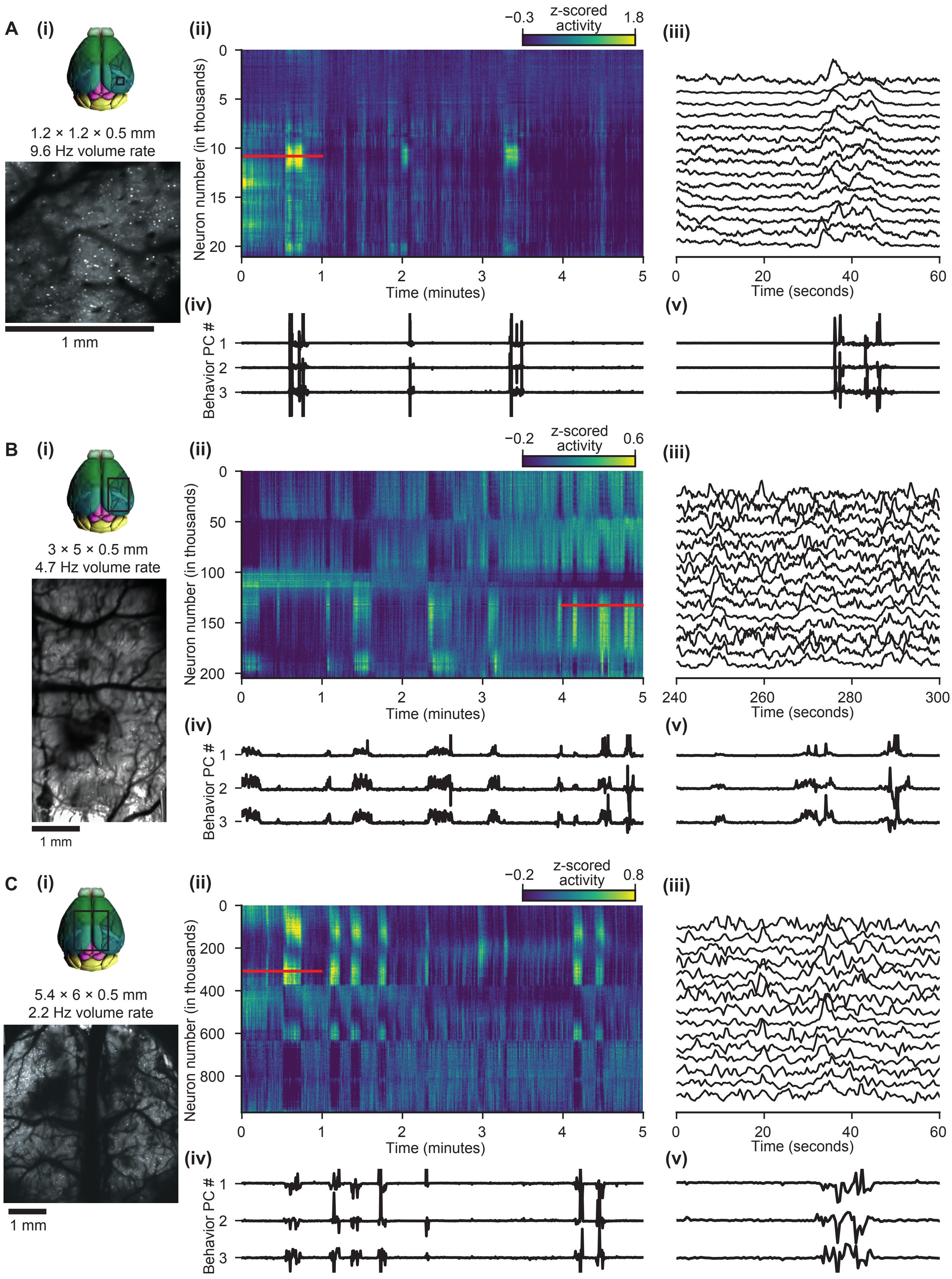

**Figure S1, continued**: related to Figure 1. Example recording configurations used in this study

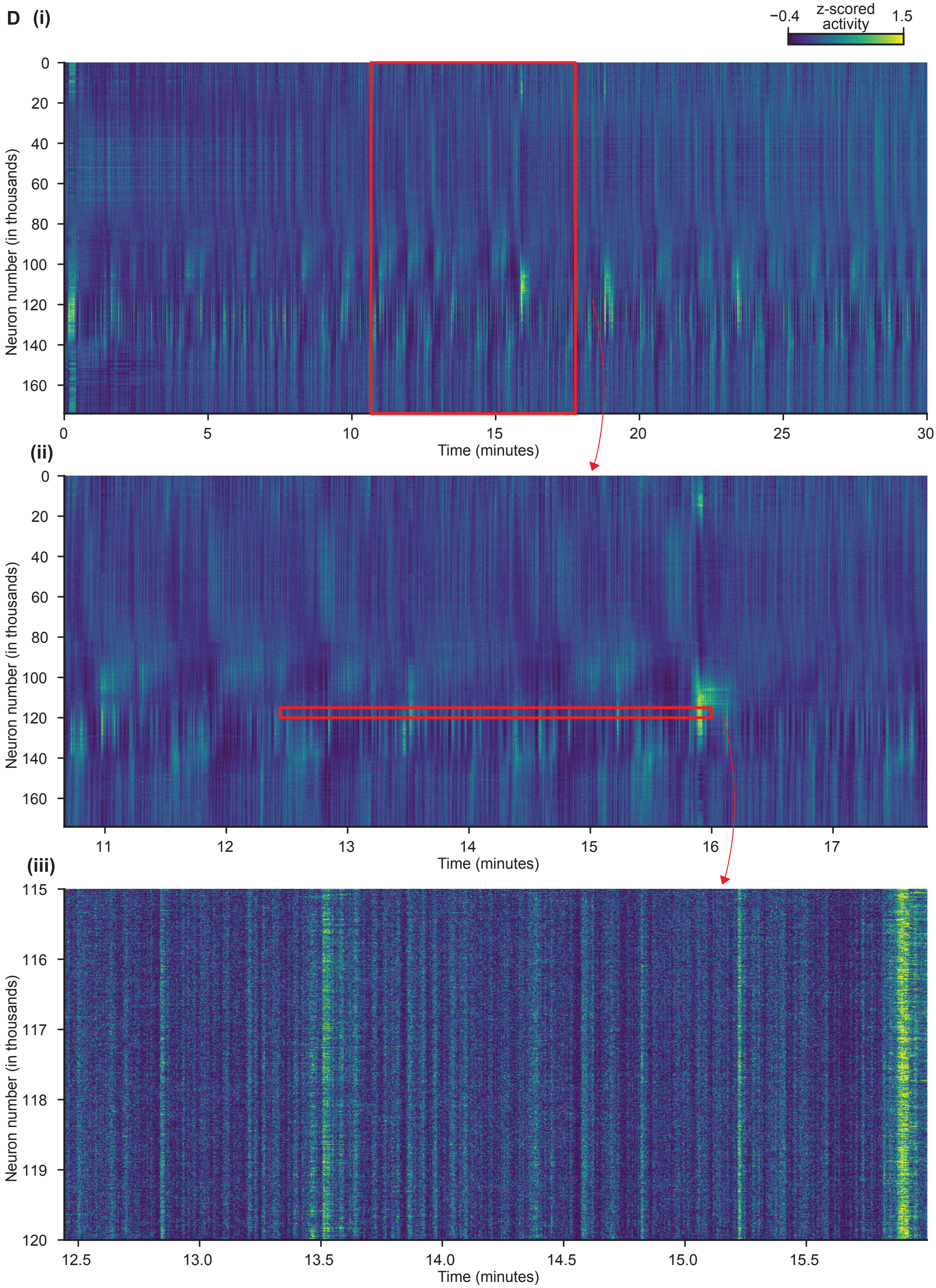
