## Supplementary figures and images for "Simultaneous, cortex-wide and cellular-resolution neuronal population dynamics reveal an unbounded scaling of dimensionality with neuron number"

### Supplemental Figure 2

**Figure S2**: related to Figure 2. Additional neural SVCA results

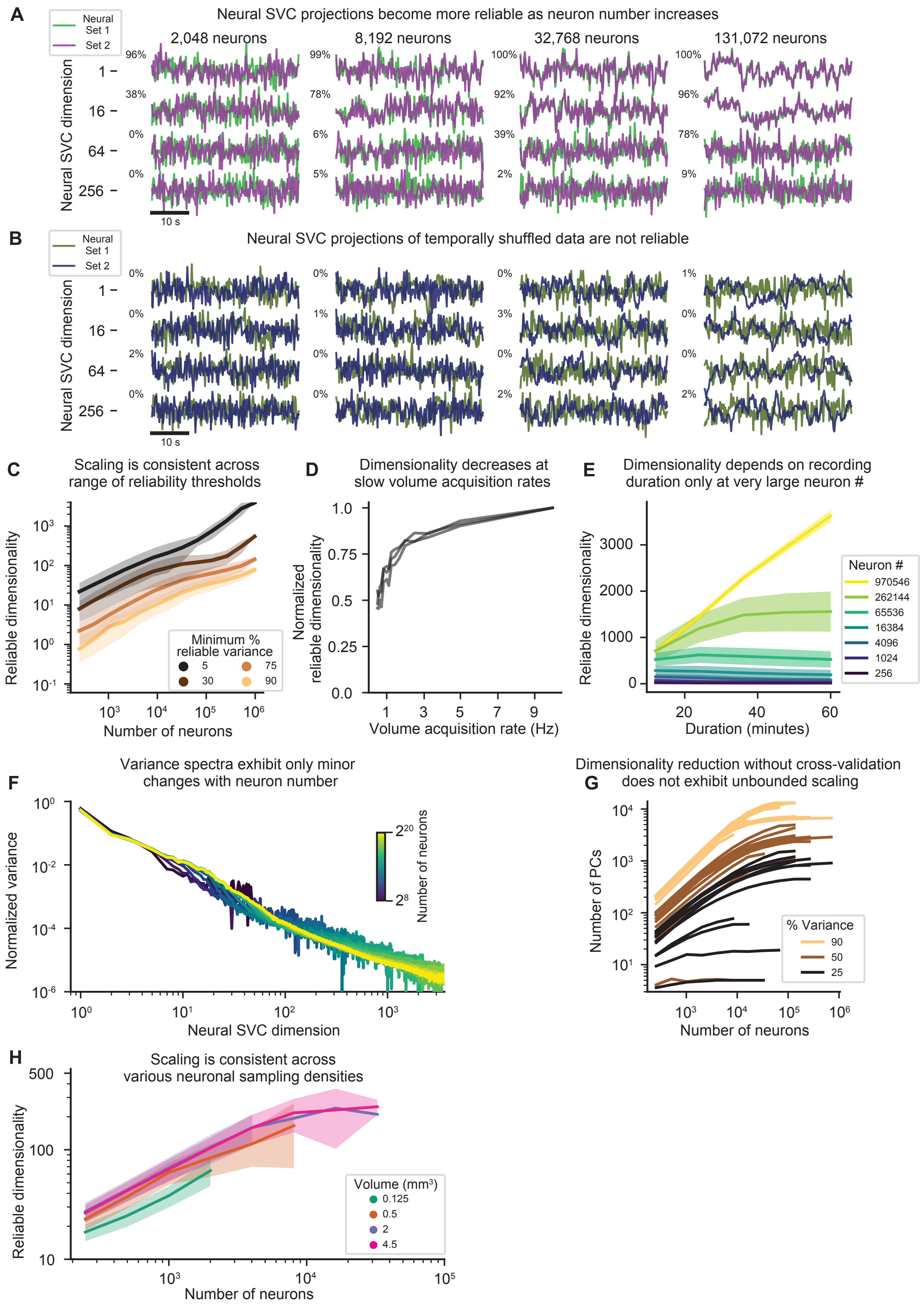

### Supplemental Figure 4

**Figure S4**: related to Figures 4 & 5. Additional neural SVC spatiotemporal characteristics

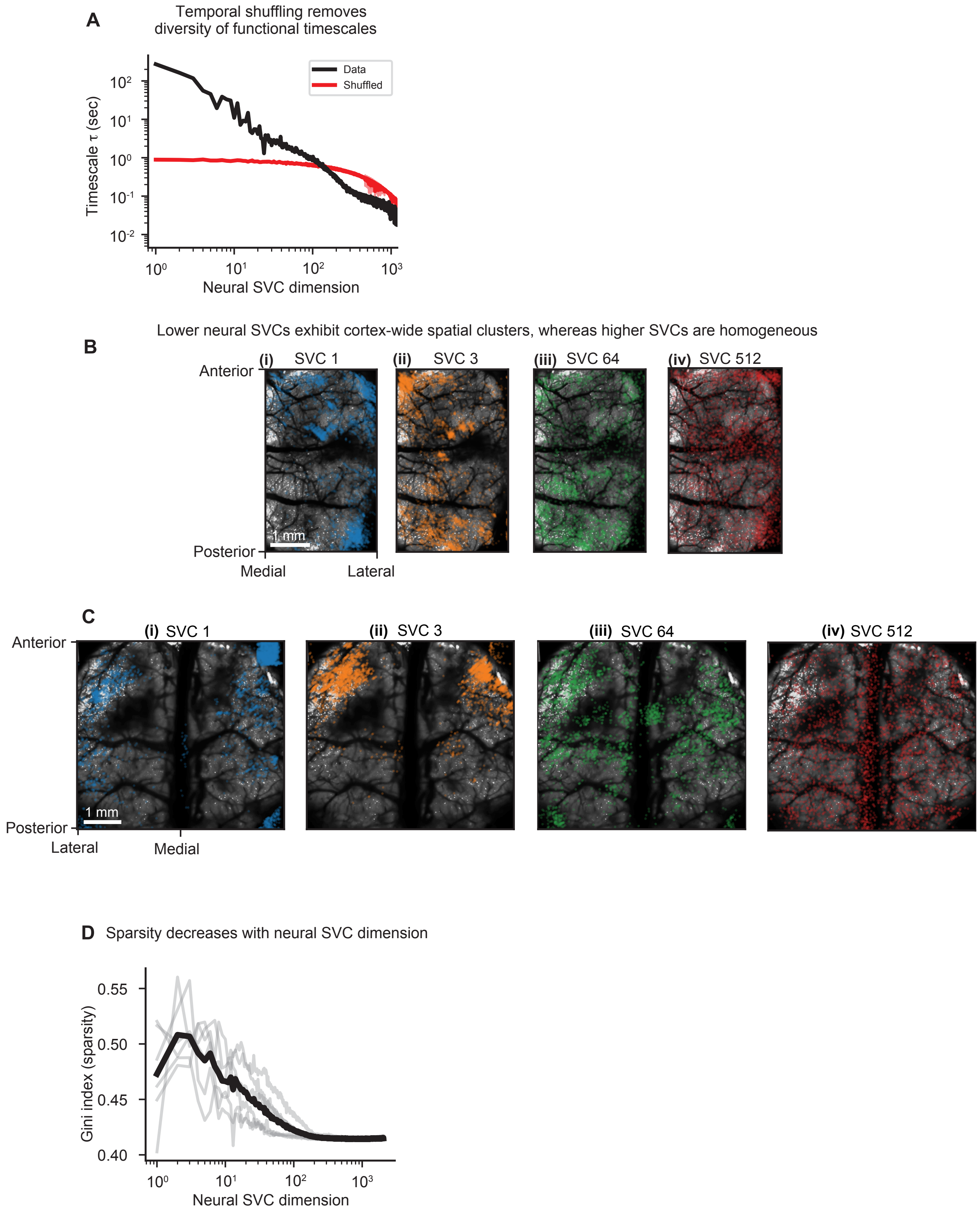
