## Supplemental Figure 3 for "Simultaneous, cortex-wide and cellular-resolution neuronal population dynamics reveal an unbounded scaling of dimensionality with neuron number"

**Figure S3**: related to Figure 3. Quantification of behavior and prediction of neural SVCs from multi-timepoint behavior

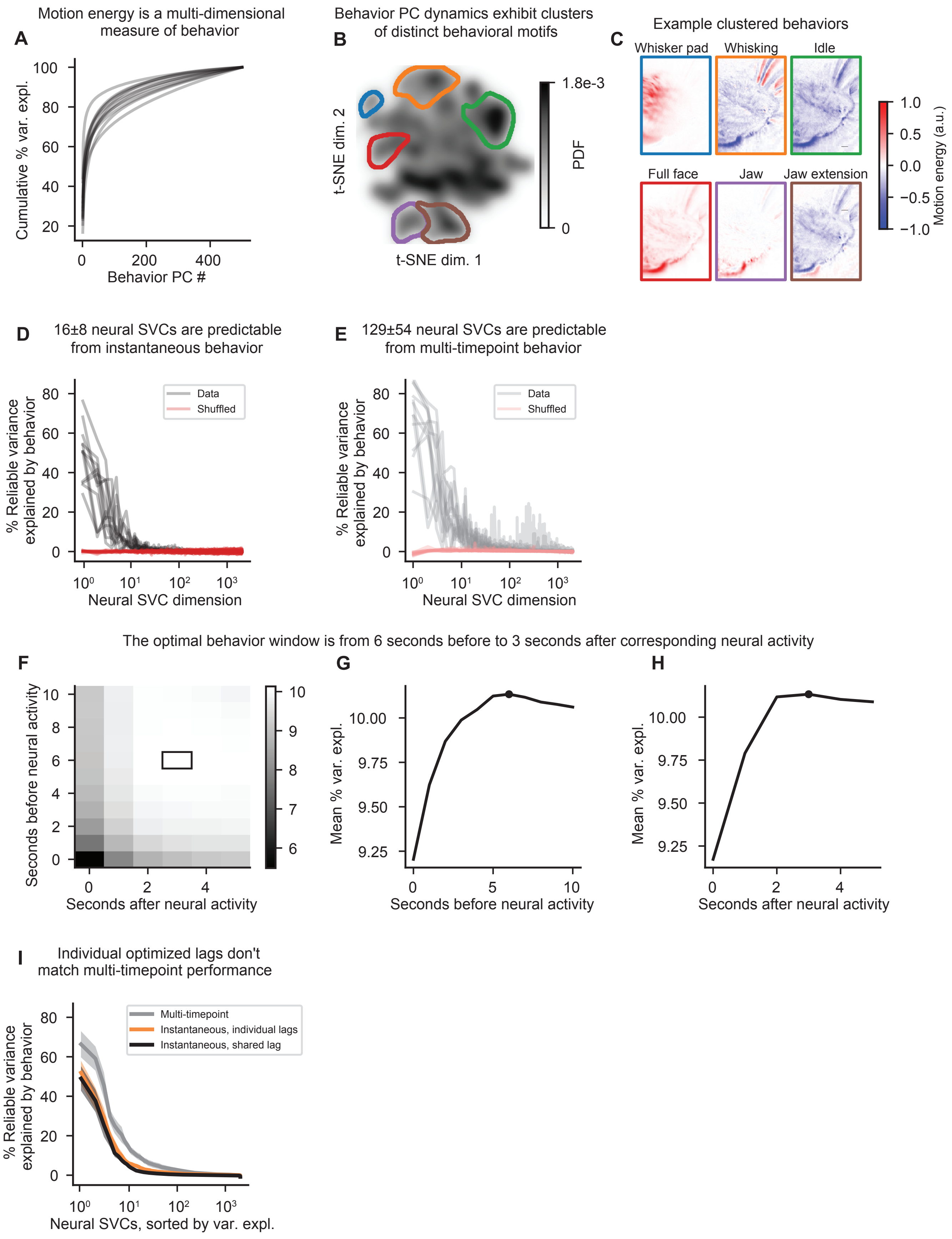
